# KDM2B regulates hippocampal morphogenesis by transcriptionally silencing Wnt signaling in neural progenitors

**DOI:** 10.1101/2023.01.26.525636

**Authors:** Bo Zhang, Chen Zhao, Wenchen Shen, Wei Li, Yue Zheng, Xiangfei Kong, Junbao Wang, Xudong Wu, Ying Liu, Yan Zhou

**Affiliations:** Department of Neurosurgery, Medical Research Institute, Frontier Science Center of Immunology and Metabolism, Zhongnan Hospital of Wuhan University, Wuhan University, Wuhan, China; Department of Cell Biology, Tianjin Medical University; Department of Neurosurgery, Tianjin Medical University General Hospital; Tianjin, China

## Abstract

The hippocampus plays major roles in learning and memory. Similar to other parts of the brain, the development of hippocampus requires precise coordination of patterning, cell proliferation, differentiation, and migration, with both cell-intrinsic and extrinsic mechanisms involved. Here we genetically removed the chromatin-association capability of KDM2B - a key component of the variant Polycomb repressive complex 1 (PRC1) - in the progenitors of developing dorsal telencephalon (*Kdm2b^ΔCxxC^*) to surprisingly discover that the size of *Kdm2b^ΔCxxC^* hippocampus, particularly the dentate gyrus, became drastically smaller with disorganized cellular components and structure. *Kdm2b^ΔCxxC^* mice displayed prominent defects in spatial memory, motor learning and fear conditioning. The differentiation trajectory of the developing *Kdm2b^ΔCxxC^* hippocampus was greatly delayed, with a significant amount of TBR2-expressing intermediate progenitors stuck along the migratory/differentiation path. Transcriptome and chromatin immunoprecipitation studies of neonatal hippocampi and their progenitors indicated that genes implicated in stemness maintenance, especially components of canonical Wnt signaling, could not be properly silenced by PRC1 and PRC2. Activating the Wnt signaling disturbed hippocampal neurogenesis, recapitulating the effect of KDM2B loss. Together, we unveiled a previously unappreciated gene repressive program mediated by KDM2B that controls progressive fate specifications and cell migration, hence morphogenesis of hippocampus during development.

**Graphic Abstract:** Loss of KDM2B-CxxC reduces repressive histone modifications – H2AK119ub and H3K27me3 - on key Wnt signal genes, hence leading to the block of their attenuation over time. Hampered differentiation and migration of hippocampal progenitors leads to hippocampal agenesis.

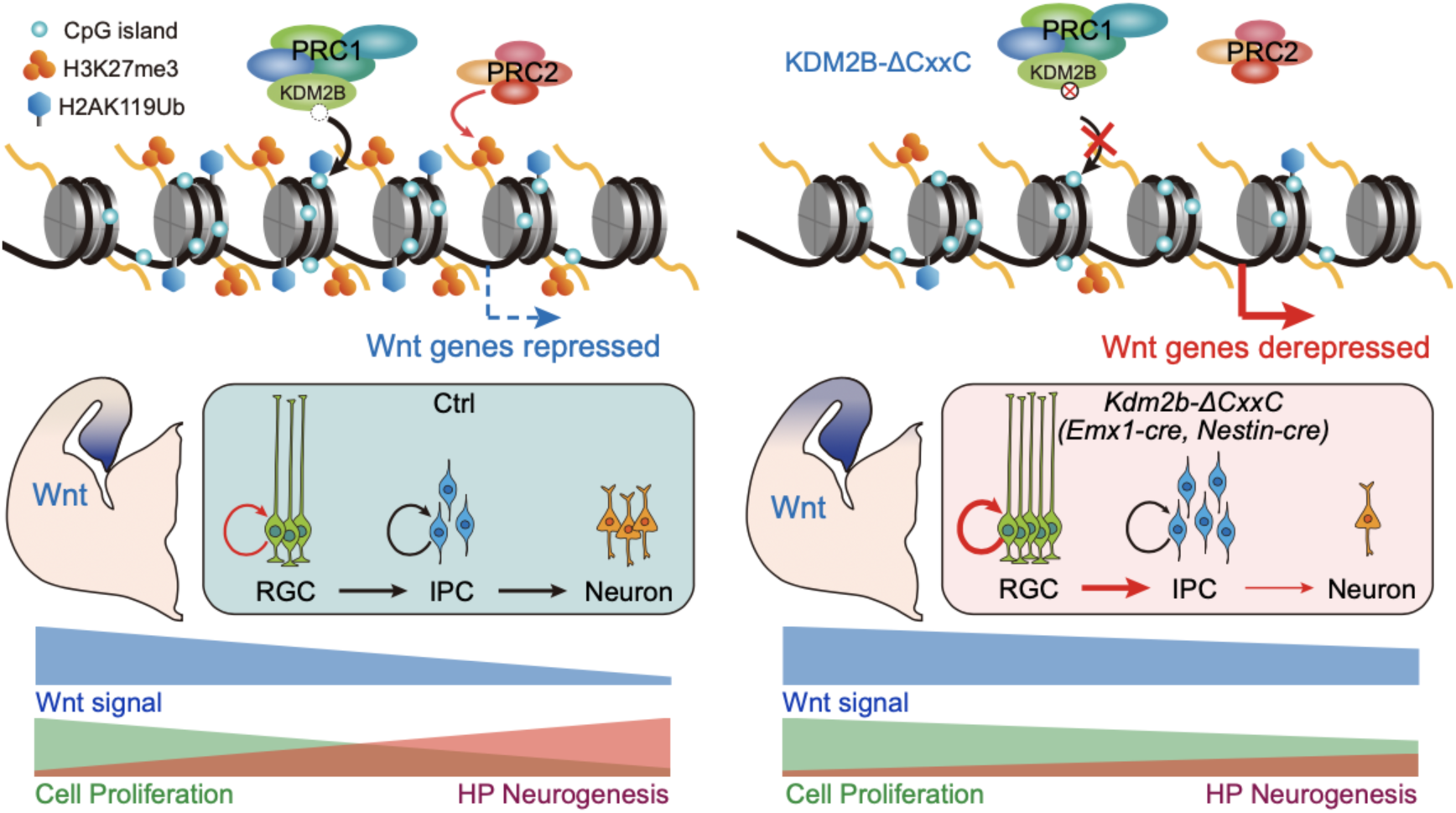

## Introduction

The hippocampus of the mammalian brain includes three major compartments: the hippocampus proper, which can be divided into three pyramidal subregions [cornu ammonis (CA) fields], the dentate gyrus (DG), and the subiculum. The hippocampus plays essential roles in spatial memory that enables navigation, in the formation of new memories, as well as in regulating mood and emotions(Anacker & Hen, 2017). Constructing a functional hippocampus requires precise production, migration, and assembly of a variety of distinct cell types during embryonic and early postnatal stages. Largely resembling neurogenesis of neocortical pyramidal neurons, the production of pyramidal neurons in CAs and granule cells in dentate gyrus follows the path of indirect neurogenesis(G. Li & Pleasure, 2014), i.e., PAX6-expressing radial glial progenitor cells (RGCs) first give rise to TBR2^+^ intermediate progenitor cells (IPCs), which then produce NeuN^+^ neurons. Particularly, the formation of the dentate gyrus relies on sequential emergence of germinative foci at different locations, including the dentate notch around embryonic (E) day 13.5 in mice, the fimbriodentate junction (FDJ) around E15.5 and the hilus around birth, which generate granule cells at different parts of DG(G. Li & Pleasure, 2014; Nelson et al., 2020). Cell fate defects and skewed cell migration during hippocampal development underly a cohort of human neurologic and psychiatric diseases(Zhong et al., 2020). Moreover, the subgranule zone (SGZ) of adult DG contains neural stem cells (NSCs) to support neurogenesis, which might have implications in the formation of new memory.

The cell fate specification during development requires coordinated actions of transcription factors (TFs), epigenetic factors and *cis*-acting elements to ensure precise gene expression and silencing. Notably, repressive histone modifications including mono-ubiquitinated histone H2A at lysine 119 (H2AK119ub1) and trimethylated histone H3 at lysine 27 (H3K27me3), respectively modified by the Polycomb repressive complex 1 (PRC1) and PRC2, are essential fate regulators in embryogenesis, organogenesis and tissue homeostasis(Blackledge & Klose, 2021; Laugesen & Helin, 2014; Piunti & Shilatifard, 2016; Schuettengruber & Cavalli, 2009; von Schimmelmann et al., 2016). The PRC1 can be recruited to the chromatin *via* either one of the CBX proteins – *a.k.a.* canonical PRC1, or other adapter proteins including KDM2B – *a.k.a.* variant PRC1(Farcas et al., 2012; Fursova et al., 2019; Z. Gao et al., 2012; He et al., 2013; Wu, Johansen, & Helin, 2013). Interestingly, PRC1 and PRC2 can reciprocally recognize repressive histone modifications mediated by each other or itself to stabilize repressive chromatin environments(Blackledge et al., 2014; Kalb et al., 2014; Margueron et al., 2009; Sugishita et al., 2021; Tamburri et al., 2020). It has been reported that the PRC2 is required for hippocampal development and the maintenance of adult neural stem cell pool of the DG, but the underlying cellular processes and molecular mechanisms were large elusive(Liu et al., 2019; Zhang et al., 2014). Furthermore, very little is known to what extent and how PRC1 is involved in hippocampal development.

KDM2B, previously known as JHDM1B, FBXL10, or NDY1, can recruit PRC1 to non-methylated CpG islands (CGIs), particularly at promoter regions, *via* its CxxC Zinc finger (ZF)(Farcas et al., 2012; Gearhart, Corcoran, Wamstad, & Bardwell, 2006; He et al., 2013; Z. Wang et al., 2018; Wu et al., 2013). The long isoform of KDM2B – KDM2BLF – contains a Jmjc demethylation domain which removes the di-methylated lysine 36 of histone H3 (H3K36me2) to regulate pluripotency and early embryogenesis(Huo et al., 2022). Mutations of human *KDM2B* gene are associated with neurodevelopment defects including intellectual disability (ID), speech delay and behavioral abnormalities(Labonne et al., 2016; van Jaarsveld et al., 2022; Yokotsuka-Ishida et al., 2021). A recent study indicated that heterozygosity of *Kdm2b* in mice impaired neural stem cell self-renewal and leads to ASD/ID-like behaviors(Y. Gao et al., 2022). However, there is no in-depth analysis to dissect how PRC1-and/or demethylase-dependent roles of KDM2B participates in multiple facets of neural development, including self-renewal, migration, differentiation and localization of neural progenitors and their progeny.

We previously showed that KDM2B controls neocortical neuronal differentiation and the transcription of *Kdm2blf* is *cis*-regulated by a long non-coding RNA which is divergently transcribed from the promoter of *Kdm2blf*(W. Li et al., 2020). However, questions remain regarding to what extent and how KDM2B regulates neural development. Here we ablated the chromatin association capability of KDM2B in the developing dorsal forebrain to surprisingly find the morphogenesis of hippocampus, especially the DG, was greatly hampered. Moreover, intermediate progenitors could not properly migrate and differentiate upon dissociating KDM2B from the chromatin. The canonical Wnt signaling were aberrantly activated in mutant hippocampi, probably due to decreased enrichment of H2AK119ub and H3K27me3 in CGI promoters of key Wnt pathway genes.

## Results

### Removing KDM2B’s chromatin association capability causes agenesis of the hippocampus

KDM2B has two main isoforms: the long isoform KDM2BLF contains a demethylase Jmjc domain while both isoforms share the CxxC zinc finger (ZF), the PHD domain, a F-box and the LRR domain. We generated the conditional knockout allele of *Kdm2b* by flanking exon 13 that encodes the CxxC ZF with two *loxP* sequences (**Figure S1A**). These floxed *Kdm2b* mice, *Kdm2b^flox(CxxC)^*, were crossed with *Emx1*-Cre and *Nestin*-Cre mice to generate conditional *Kdm2b^Emx1-ΔCxxC^* and *Kdm2b^Nestin-ΔCxxC^* conditional knockout (cKO) mice respectively to abolish KDM2B’s association with the chromatin. Although *Kdm2b^Nestin-ΔCxxC^* mice could not survive past postnatal day 7 (P7), *Kdm2b^Emx1-ΔCxxC^* mice were born at the mendelian ratio and thrive through adulthood without gross abnormality (**Figure S1B-S1C**). Quantitative RT-PCR experiments confirmed the deletion of exon 13, while transcriptions of other parts of *Kdm2b*, including the region encoding the C-terminal LRR domain, remained intact in cKO neocortices (**Figure S1D-S1F**). We then performed *in situ* hybridization of neonatal (postnatal day 0, P0) brains using probes targeting exon 13, which showed *Kdm2b* is expressed in the developing hippocampus across the CA region and the DG of control *Kdm2b^flox/flox^* brains (**Figure S1G**). Importantly, the expression of *Kdm2b-CxxC* was almost gone in the *Kdm2b^Emx1-ΔCxxC^* hippocampi (**Figure S1H**). Moreover, immunoblotting of embryonic day 15.5 *Kdm2b^Nestin-ΔCxxC^* neocortex revealed truncated long and short isoforms of KDM2B, confirming the selective deletion of the CxxC ZF, which could abolish the CGI association by KDM2B and variant PRC1.1 (**Figure S1I**).

Strikingly, the hippocampi, particularly the DGs, of adult *Kdm2b^Emx1-ΔCxxC^* cKO brains were greatly shrunk in size in all examined sections (**Figure 1A, S2B**). Although Wfs1-expressing pyramidal cells could be detected in the CA1 region of *Kdm2b^Emx1-ΔCxxC^* cKO brains (**Figure S2A**), Calbindin-expressing cells were mostly diminished (**Figure 1B**). Both upper and lower blades of cKO DGs were much shorter than controls (**Figure 1C; S1C**), with numbers of NeuN+ granule cells and HopX+ NSCs decreased by 39.7% and 56.8% respectively (**Figure 1D-1G**). Immuno-staining showed irregularity and ectopic dispersion of Calbindin+, ZBTB20+ or NeuN+ granule cells in cKO DGs (**Figure 1B-1D**), accompanied with fewer SOX2+ or GFAP+ NSCs, fewer TBR2+ neuroblasts, and fewer PROX1+ or DCX+ neurons in the cKO SGZ (**Figure 1H; S1C**). Interestingly, many HOPX+ NSCs were found to be ectopically localized inside the granule cell layer of cKO DGs (**Figure 1F**, red arrows). The ventricles were also enlarged in cKO brains along with thinner neocortices (**Figure 1A, S2D**). The enlarged ventricles of cKO brains could already be seen at P0 for unknown reasons without thinning of neocortices (**Figure S2G-S2K**). At P7, numbers of upper-layer (SATB2+) and lower-layer (CTIP2+) neocortical neurons were not significantly altered upon CxxC deletion of KDM2B (**Figure S2L-S2M**). Consistently, numbers of PAX6-expressing radial glial progenitors and TBR2-expressing intermediate progenitors were not changed in E16.5 cKO neocortices (data not shown). Together, ablation of the chromatin association capability of KDM2B in developing dorsal forebrains causes hippocampal agenesis, while the thinning of adult cKO neocortices could be secondary to ventricle dilation.

**Figure 1.**
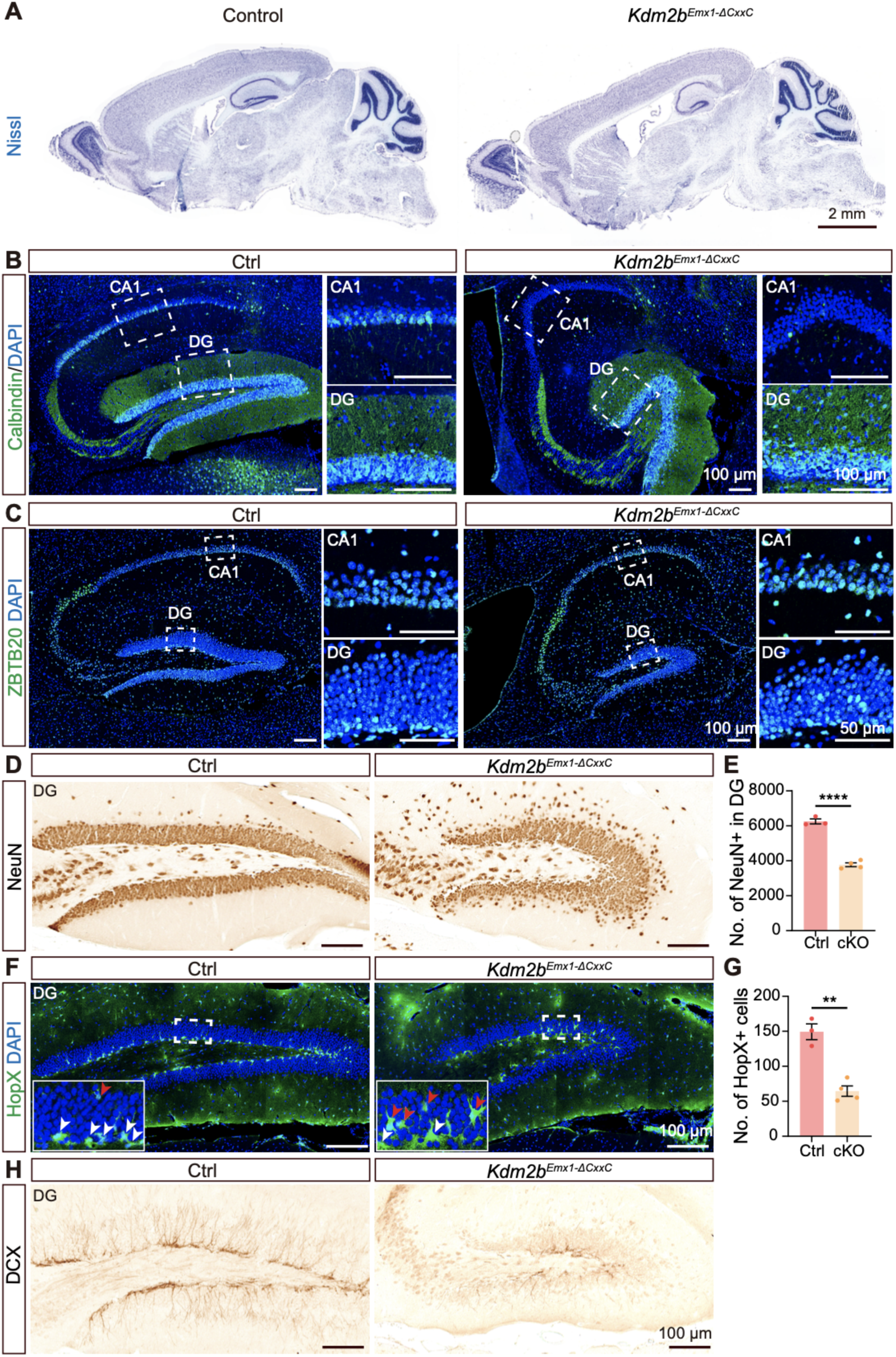
Deletion of the KDM2B-CxxC causes agenesis of hippocampus. (A) Representative images showing Nissl staining on sagittal sections of adult control and *Kdm2b^Emx1-ΔCxxC^* brains. (B, C) Immunofluorescent (IF) staining of Calbindin (B) and ZBTB20 (C) on sagittal sections of adult control (left) and *Kdm2b^Emx1-ΔCxxC^* (right) hippocampi. Nuclei were labeled with DAPI (blue). Boxed CA1 and dentate gyri (DG) were enlarged on the right. (D, H) Immunohistochemical (IHC) staining of NeuN (D) and DCX (H) on sagittal sections of adult control (left) and *Kdm2b^Emx1-ΔCxxC^* (right) dentate gyri (DG). (F) IF staining of HopX on sagittal sections of adult control (left) and *Kdm2b^Emx1-ΔCxxC^* (right) DG. Boxed regions were enlarged on bottom-left corners. White and red arrows denote HopX+ signals in the subgranule zone (SGZ) and granule cell layer respectively. (E, G) Quantification of NeuN+ (E) and HopX+ (G) cells in the DG. n = 3 for control brains and n = 4 for *Kdm2b^Emx1-ΔCxxC^* brains. Data are represented as means ± SEM. Statistical significance was determined using an unpaired two-tailed Student’s t-test (E, G). **P<0.01; ****P<0.0001. Scale bars, 2mm (A), 100 μm (B-D, F, H), 50 μm (C CA1 and DG). DG, dentate gyrus.

### *Kdm2b^Emx1-ΔCxxC^* mice displays defects in spatial memory, motor learning and fear conditioning

Since hippocampus is essential for spatial navigation and memory consolidation, as well as for anxiety and depression behaviors, we conducted a series of behavior tests. First, open field tests showed reluctance of *Kdm2b^Emx1-ΔCxxC^* mice to explore center regions of open fields (**Figure 2A-2B**), which could be decreased willingness of cKO mice to explore and/or increased anxiety. The distance and velocity traveled, and immobility time in open field tests were not significantly altered in *Kdm2b^Emx1-ΔCxxC^* mice (**Figure S3A**). The immobility time of *Kdm2b^Emx1-ΔCxxC^* mice in forced swimming tests were a tad shorter than control mice (no statistical significance), whereas the immobility time of cKO mice in tail suspension test was the same as controls, suggesting cKO mice did not display depression-related behavior (**Figure S3B-S3C**). Interestingly, cKO mice spent shorter time in open arms of elevated plus maze (no statistical significance), indicating that cKO mice tend to be hypersensitive to anxiety (**Figure S3D**). Secondly, the rotarod performance tests indicated that *Kdm2b^Emx1-ΔCxxC^* cKO mice have compromised capacity for motor coordination and learning, as they endured shorter time in rotarods than controls (**Figure 2C**). Third, Morris water maze tests revealed that *Kdm2b^Emx1-ΔCxxC^* cKO mice were defective in spatial learning. It took cKO mice longer time to find the platform in the training stage (**Figure 2D**). Consistently, in the probe trial, *Kdm2b^Emx1-ΔCxxC^* cKO mice spent shorter time in quadrant holding the platform (**Figure 2I**), whereas control and cKO mice showed no differences in swimming distances and velocity (**Figure 2E-2H**). Fourth and most importantly, in contrast to control mice, *Kdm2b^Emx1-ΔCxxC^* cKO mice failed to display prolonged freezing time in both context-and sound-induced fear conditioning tests (**Figure 2J-2M**). Together, ablation of the chromatin association capability of KDM2B leads to development failure of the hippocampus, which might cause defects in spatial memory, motor learning and fear conditioning.

**Figure 2.**
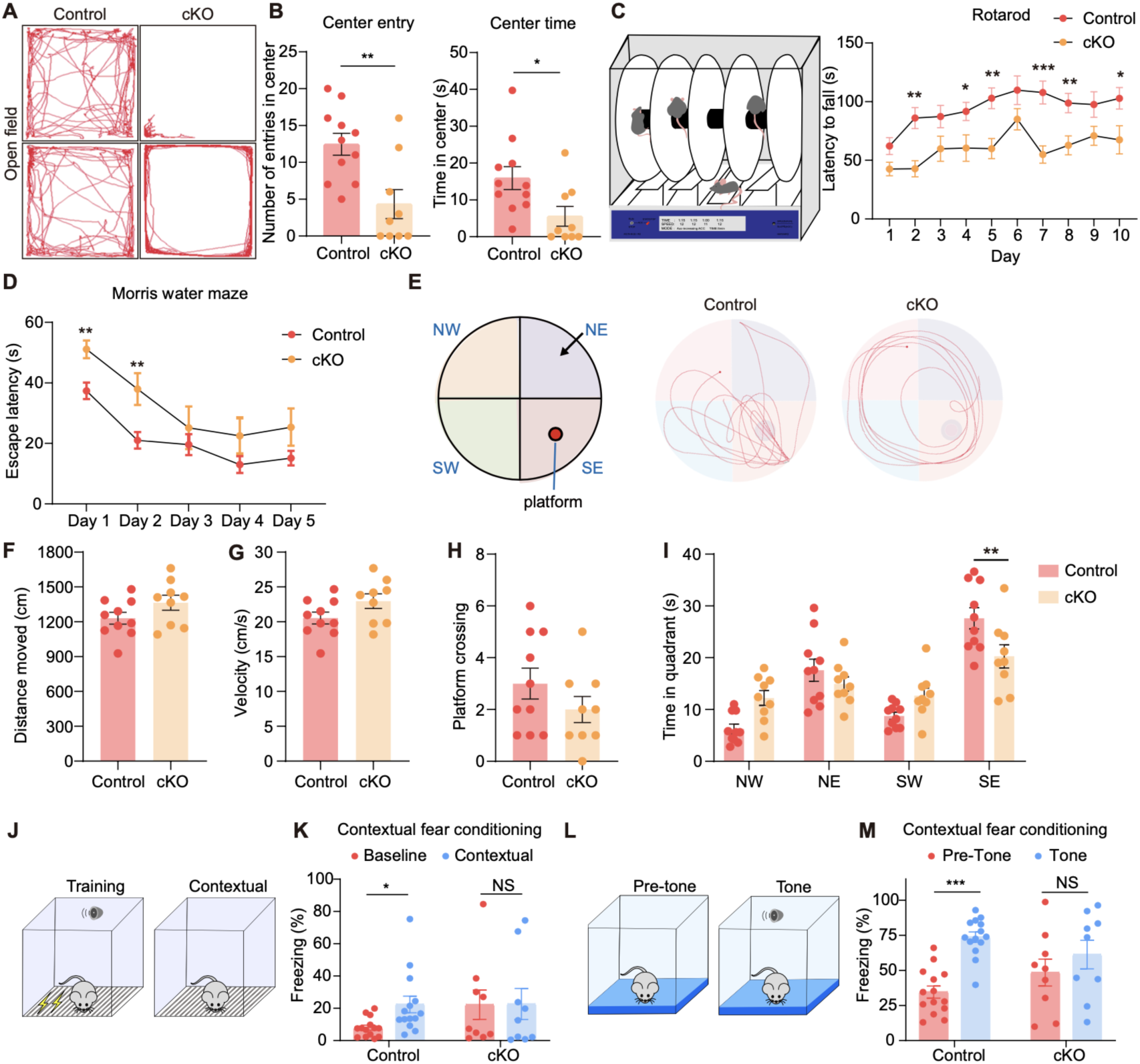
*Kdm2b^Emx1-ΔCxxC^* mice exhibit defects in motor learning, spatial memory, and contextual fear conditioning. (A) Representative traces of control mice and *Kdm2b^Emx1-ΔCxxC^* mice in the open-field arena. (B) Quantification of number of entries in center, and time spent in the center in the open-field test. (C) Latency to fall during the rotarod test. (D) Latency to find the hidden platform across training period in the Morris water maze test. (E) An overhead view of the Morris water maze, and representative swim paths of control mice and *Kdm2b^Emx1-ΔCxxC^* mice during the probe trial. The platform was set in the SE quadrant. (F, G) Distance moved (F) and velocity (G) during the probe trial (platform removed). (H) Frequencies of platform crossing during the probe trial. (I) Time spent in each quadrant during the probe trial. (J, K) The proportion of freezing time in context before training (Baseline) and after training (Contextual). (L, M) The proportion of freezing time in a new context before tone (Pre-Tone) and after tone (Tone). Data are represented as means ± SEM. Statistical significance was determined using two-way ANOVA followed by Sidak’s multiple comparisons test (E, M), or using an unpaired two-tailed Student’s t-test (F, H-K). *P < 0.05, **P < 0.01, ***P < 0.001, and ****P < 0.0001; NS, not significant. n = 11 mice in (A-C), n = 10 in (D-I), n = 14 in (J-M) for control and n = 10 mice for *Kdm2b^Emx1-ΔCxxC^*.

### Deletion of the CxxC ZF of KDM2B has no effect on adult neurogenesis of the DG

The shrunken hippocampi and diminished NSC pool in *Kdm2b^Emx1-ΔCxxC^* cKO mice prompted us to investigate whether the ablation of KDM2B’s chromatin association capability could hamper adult neurogenesis of the DG. We thus crossed the *Kdm2b^flox(CxxC)^* mice with *Nestin-CreERT2* mice to produce *_Kdm2bNestin-CreERT2-ΔCxxC_* _cKO mice. Adult *Kdm2b*_*_Nestin-CreERT2-ΔCxxC_* _cKO and_ control mice were administered with tamoxifen (TAM) for six consecutive days to ablate the CxxC ZF in adult NSCs and their progeny. BrdU was also administered for six consecutive days to label NSCs’ progeny (**Figure S3E**). Brains were collected at day 8 and day 35 of post-TAM injection for immunofluorescent staining of BrdU, along with DCX (day 8) or PROX1 (day 35), two markers for new-born and mature granule neurons respectively. Data showed numbers of BrdU-labelled cells and DCX^+^BrdU^+^ double-positive cells, and ratios for DCX^+^BrdU^+^/BrdU^+^ cell were comparable between *Kdm2b^Nestin-CreERT2-ΔCxxC^* cKO and control DGs in all examined sections at day 8 (**Figure S3F-S3G**). Similarly, at day 35, numbers of BrdU-labelled cells and PROX1^+^BrdU^+^ double-positive cells, and ratios for PROX1^+^BrdU^+^/BrdU^+^ cell were not altered in *Kdm2b^Nestin-CreERT2-ΔCxxC^* cKO DGs (**Figure S3H-S3J**). Thus, removal of KDM2B’s chromatin association capability in adult NSCs exerts no effect on adult neurogenesis of DGs.

### Hampered migration of intermediate progenitors and neurogenesis of granule cells upon loss of KDM2B

In DG development, neural progenitors, especially TBR2-expressing intermediate progenitor cells (IPCs) migrate from the dentate neuroepithelial (DNe) stem zone (the primary - 1ry matrix) through the dentate migratory stream (DMS, the secondary - 2ry matrix) to the developing DG (the 3ry matrix), while being distributed in multiple transient niches (**Figure 3D**). The prominent DG defects in *Kdm2b^Emx1-ΔCxxC^* cKO brains prompted us to examine distributions of intermediate progenitors and neurons along the migratory path. P0 brain sections were co-stained with TBR2 to label IPCs and with GFAP to label astrocytic scaffold at the fimbriodentate junction (FDJ) of the DMS and the fimbria. Total number of IPCs were increased by 15.3% upon loss of KDM2B-CxxC. Strikingly, in *Kdm2b^Emx1-ΔCxxC^* cKO brains, significant more TBR2+ IPCs were accumulated at the DNe and the DMS, along with significant fewer TBR2+ IPCs at the DG (**Figure 3A-3E**). Of note, TBR2+ IPCs were more loosely distributed at the FDJ, with GFAP-labelled astrocytic scaffold scattered at FDJ but constricted at fimbria (**Figure 3A-3C**). We next carried out EdU birthdating experiments by administering EdU at E15.5 followed by EdU and PROX1 co-labelling at P2. Consistently, 31.3% fewer PROX1-labelled granule cells could be detected in cKO DGs but 259.0% more in the cKO FDJ. In cKO brains, 33.3% fewer E15.5-labelled EdU cells reached the DG and co-stained with PROX1, while ectopic PROX1 cells were detected at FDJ (**Figure 3F-3H**). Furthermore, proportions of PROX1+ cells in lower and upper blades of DGs were switched into a lower-more and upper-fewer status in cKO DGs (**Figure 3I-3K**). Congruently, by P7, a big chunk of PROX1+ cells could be seen at the FDJ of *Kdm2b^Emx1-ΔCxxC^* brains (**Figure S4F**). Together, loss of KDM2B-CxxC impedes migration and differentiation of IPCs, hence proper production and localization of granule neurons during hippocampal formation.

**Figure 3.**
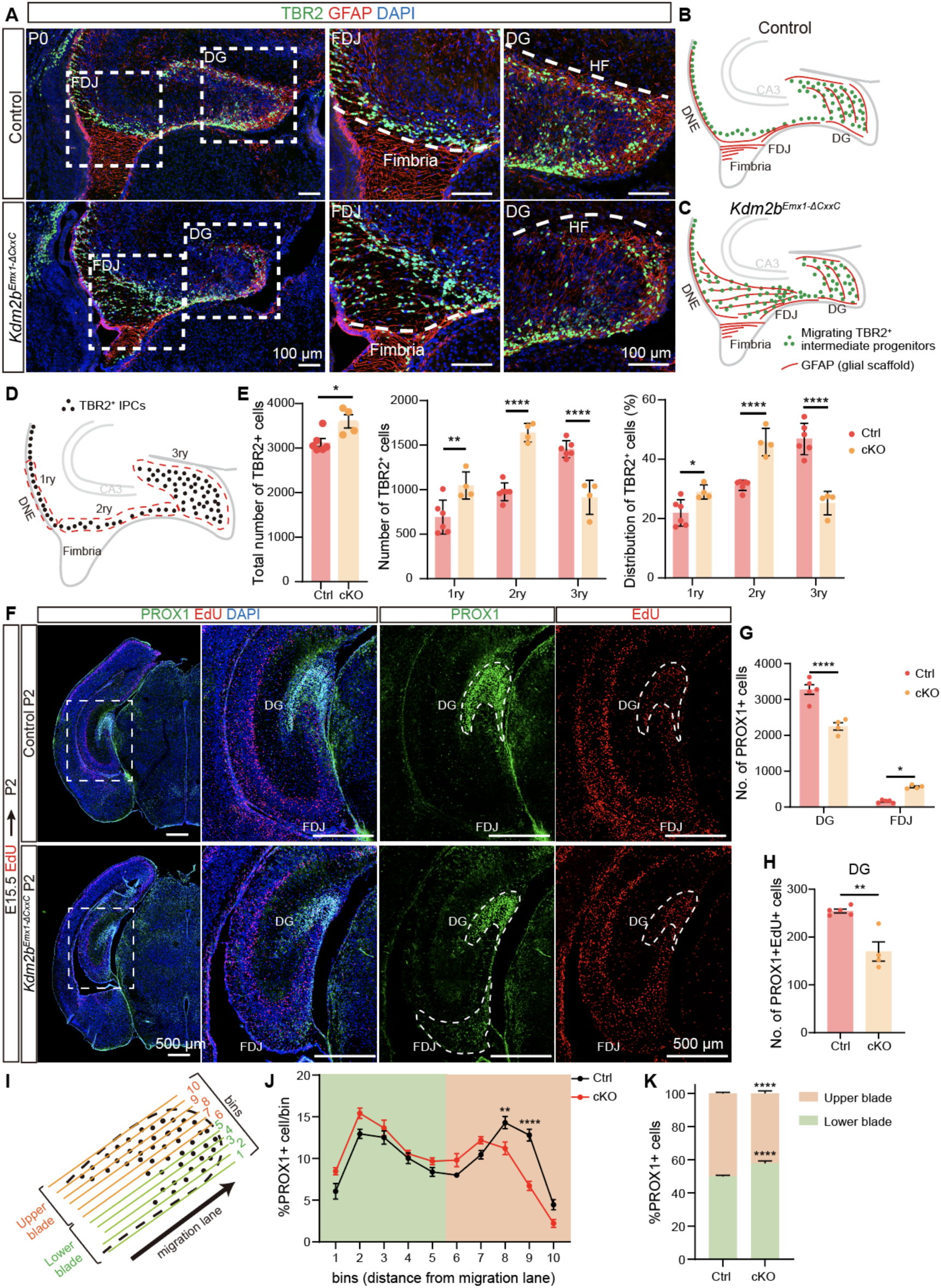
Ablation of the KDM2B-CxxC impedes the migration of intermediate progenitors and production of granular cells. (A) Double immunofluorescence of TBR2 (green) and GFAP (red) on P0 wild-type and *Kdm2b^Emx1-ΔCxxC^* hippocampi. Nuclei were labeled with DAPI (blue). Boxed regions of FDJ and DG were enlarged on the right. (B, C) The schematic of P0 wild-type and *Kdm2b^Emx1-ΔCxxC^* hippocampi. Green dots represent migrating TBR2+ intermediate progenitors, and red lines represent GFAP+ glial scaffold. (D, E) Distribution of TBR2+ cells along the three matrices, where dashed lines demarcate regions considered as 1ry, 2ry, and 3ry matrix (D). n = 6 for control brains and n = 4 for *Kdm2b^Emx1-ΔCxxC^* brains. (F) EdU was administrated at E15.5 and double labeling of PROX1 and EdU was performed on P2 coronal sections. Boxed regions were enlarged on the right, and single channel fluorescence staining of PROX1 and EdU were shown respectively. Dashed lines indicate DG and FDJ. (G) Quantification of PROX1+ cells in DG and FDJ. (H) Quantification of PROX1+EdU+ cells in DG. (I) Analysis of the distribution of PROX1+ granule neurons within the upper and lower blade of the forming DG at P2: the 3ry matrix was divided into 10 ventral--to-dorsal bins spanning the lower to upper blade domain. (J) PROX1+ cells were counted within each bin. The percentage of PROX1+ cells in each bin is represented. (K) Quantification of the percentage of PROX1+ granule neurons positioned in the DG lower blade (bins 1–5), versus the DG upper blade (bins 6–10) in P2 controls and *Kdm2b^Emx1-ΔCxxC^*. n = 5 for control brains and n = 4 for *Kdm2b^Emx1-ΔCxxC^* brains (F-K). Data are represented as means ± SEM. Statistical significance was determined using an unpaired two-tailed Student’s t-test (E left, H), or using two-way ANOVA followed by Sidak’s multiple comparisons test (E middle and right, G, J, K). *P < 0.05, **P < 0.01, ***P < 0.001, and ****P < 0.0001. Scale bars, 100 μm (A), 500 μm (F). DG, dentate gyrus; DMS, dentate migratory stream; FDJ, fimbriodentate junction; HF, hippocampal fissure; 1ry, primary matrix; 2ry, secondary matrix; 3ry, tertiary matrix.

The hampered migration of IPCs upon loss of KDM2B-CxxC could be due to defects of the cortical hem (CH) derived astrocytic scaffolds at the fimbria. To exclude the possibility, *Kdm2b^Nestin-ΔCxxC^* cKO brains were inspected, because *Nestin* is mostly not expressed in CH derived astrocytic scaffolds(Caramello, Galichet, Rizzoti, & Lovell-Badge, 2021). Data showed that P0 *Kdm2b^Nestin-ΔCxxC^* brains displayed almost the same phenotypes as those in *Kdm2b^Emx1-ΔCxxC^* brains, *i.e.*, overproduction of TBR2+ IPCs, significant more IPCs accumulated at the DNe and the FDJ, but fewer IPCs at the DG, suggesting that the migrating defects of IPCs upon loss of KDM2B-CxxC were largely cell-intrinsic (**Figure S4A-S4E**).

### Disturbed differentiation of neural progenitors on loss of KDM2B-CxxC

We next investigated how the RGC-IPC-Neuron neurogenesis path of hippocampi were influenced on loss of KDM2B-CxxC. At E13.5, when the hippocampal primordia just emerged, the distribution of BLBP, the marker for astrocytic progenitors at CH, were unaltered in *Kdm2b^Emx1-ΔCxxC^* cKO brains (**Figure S5A-S5B**). Similarly, the expression pattern of SOX2, the marker for neocortical and hippocampal RGCs, were not changed in CH, DNe and hippocampal neuroepithelium (HNe) upon loss of KDM2B-CxxC (**Figure S5A**, the middle panel). In addition, the distribution of PAX6 and TBR2, markers for RGC and IPCs respectively, at E13.5 were almost the same between cKOs and controls (**Figure S5A**, the bottom panel). We further examined whether the capacity of proliferation and differentiation of hippocampal progenitors was affected at E14.5 by co-labeling brain sections with PAX6, TBR2 and EdU (2 hours pulse). Data showed that deletion of KDM2B-CxxC had no effect on abundance and proliferation of PAX6+ RGCs and TBR2+ IPCs in DNEs and CHs. Consistently, the differentiation capability from RGCs to IPCs was unchanged in cKO brains, because the number of PAX6+TBR2+ cells and the ratio of PAX6+TBR2+ cells among all PAX6+ cells were comparable between cKOs and controls (**Figure S5C-S5D**). Thus, the hippocampal agenesis in *Kdm2b^Emx1-ΔCxxC^* cKO brains was not due to specification, maintenance, and differentiation of early neural progenitors.

We then moved onto E16.5, when migration and neurogenesis of hippocampal progenitors are at the climax. In control brains, most PAX6+ RGCs were localized at the DNe, while TBR2+ IPCs were more evenly distributed along the DNe-FDJ-DG migratory/differentiating path (**Figure 4A**-**4D**). Although total numbers of PAX6+ RGCs and TBR2+ IPCs were not significantly altered in cKO hippocampi (**Figure 4E**-**4F**), their distribution along the DNe-FDJ-DG path were dramatically delayed. The cKO DNes were significantly enriched with more PAX6+, PAX6+EdU+ (dividing) and TBR2+ cells (**Figure 4G**-**4I**). In contrast, the distribution of total and dividing PAX6+ and TBR2+ cells in cKO FDJs and DGs was drastically reduced, except for that of TBR2+ cells in the FDJ (**Figure 4G**-**4L**). Furthermore, the differentiating rate from RGCs to IPCs (PAX6+TBR2+/PAX6+) at FDJ is 60% higher in cKOs (**Figure 4N**), but PAX6+TBR2+ cells were 73.9% fewer in cKO DGs (**Figure 4M**). By P7, numbers of total and dividing TBR2+ IPCs were significantly decreased at cKO DGs, whereas those at hippocampal SVZ and FDJ were greatly increased (**Figure 4O**-**4S; S6A**). In summary, the migratory and differentiating trajectory of hippocampal progenitors were greatly delayed upon loss of KDM2B-CxxC.

**Figure 4.**
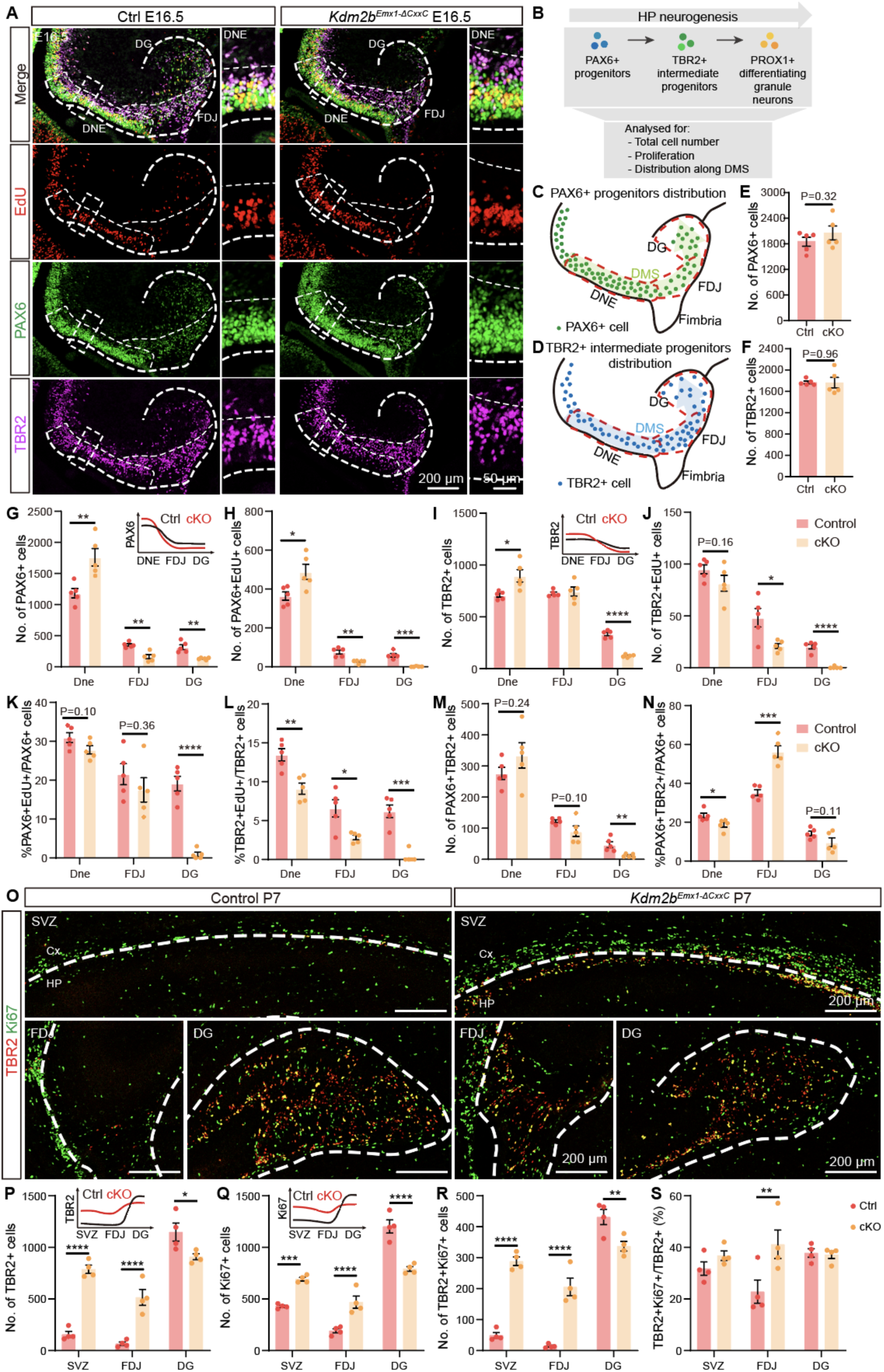
Blocked differentiation of neural progenitors on loss of KDM2B-CxxC. (A) Triple-labeling of PAX6 (green), TBR2 (violet) and EdU (red) on E16.5 control and *Kdm2b^Emx1-ΔCxxC^* brain sections. Boxed regions were enlarged on the right, and single channel fluorescence staining of PAX6, TBR2 and EdU were shown respectively. Dashed lines outline the hippocampi and distinguish DNE, FDJ and DG. (B) Experimental analysis scheme: total number of cells, the proportion of proliferative cells, and the distribution of cells along the DMS were quantified. (C, D) The schematic of E16.5 wild-type hippocampi. Green dots represent PAX6+ progenitors and blue dots represent migrating TBR2+ intermediate progenitors. Red dashed lines distinguish DNE, FDJ and DG. (E, F) Quantification of PAX6+ and TBR2+ cells. (G-N) Quantification of the distribution of PAX6+ (G), PAX6+EdU+ (H), TBR2+ (I), TBR2+EdU+ (J) and PAX6+TBR2+ cells (M), and quantification of the proportion of PAX6+EdU+/PAX6+ (K), TBR2+EdU+/TBR2+ (L) and PAX6+TBR2+/PAX6+ (N) along DMS. The distribution patterns of PAX6+ (G) and TBR2+ (I) cells in control and cKO hippocampi were shown as line graphs on the upper right corners. n = 5 for control brains and n = 5 for *Kdm2b^Emx1-ΔCxxC^* brains. (O) Double-labeling of TBR2 (red) and Ki67 (green) on P7 coronal section of control and *Kdm2b^Emx1-ΔCxxC^* SVZ, FDJ and DG. Dashed lines outline the hippocampi. Immunofluorescence staining of whole brain sections were shown in **Fig. S6A**. (P-S) Quantification of the distribution of TBR2+ (P), Ki67+ (Q) and TBR2+Ki67+ cells (R), and the proportion of TBR2+Ki67+/TBR2+ (S) at SVZ, FDJ and DG. n = 4 for control brains and n = 4 for *Kdm2b^Emx1-ΔCxxC^* brains. Data are represented as means ± SEM. Statistical significance was determined using an unpaired two-tailed Student’s t-test (E, F), or using two-way ANOVA followed by Sidak’s multiple comparisons test (G-N, P-S). *P < 0.05, **P < 0.01, ***P < 0.001, and ****P < 0.0001. Scale bars, 200 μm (A, left), 50 μm (A, right), 200 μm (O). DNE, dentate neuroepithelium; FDJ, fimbriodentate junction; DG, Dentate Gyrus; SVZ, subependymal ventricular zone; Cx, Cortex; HP, Hippocampus.

We next asked whether neuronal maturation was impeded on loss of KDM2B-CxxC. To this end, we crossed the *Kdm2b^flox(CxxC)^* mice with *Nex-Cre* mice to obtain *Kdm2b^Nex-ΔCxxC^* cKO mice, where KDM2B-CxxC was specifically ablated in all postmitotic neurons but not progenitors of neocortices and hippocampi. Phenotypic analyses revealed that hippocampal morphology and cellular components, including PROX1+ DG granule cells, of P7 *Kdm2b^Nex-ΔCxxC^* cKOs did not show any abnormality, suggesting that hippocampal agenesis was not due to defects on postmitotic neuron differentiation (**Figure S6B**). Together, KDM2B regulates hippocampal morphogenesis by controlling multiple behaviors, including coordinated RGC to IPC differentiation, migration and divisions of neural progenitors (**Figure S6C**).

### Aberrantly activated Wnt signaling due to alterations of key histone modifications in *Kdm2b* mutant hippocampus

We next sought to unveil molecular events and mechanisms underlying hippocampal agenesis caused by KDM2B mutation. First, hippocampal tissue from P0 control and *Kdm2b^Emx1-ΔCxxC^* cKO brains were harvested and subjected to RNA-seq transcriptome analyses. Data showed 886 genes and 575 genes were activated and repressed respectively in *Kdm2b^Emx1-ΔCxxC^* cKO hippocampi. In line with aforementioned phenotypic analyses, expression levels of markers for progenitor cells including *Pax6*, *Neurog2* and *TBR2*/*Eomes* were increased in cKO hippocampi (**Figure 5A**). Gene ontology (GO) analyses revealed that up-regulated genes were involved in pattern formation, morphogenesis, cell proliferation and canonical Wnt signal pathways (**Figure 5B, S8A**), whereas down-regulated genes encompassing those associated with neuronal structures and functions (**Figure 5C**). Notably, a series of canonical Wnt pathway components, including ligands, receptors, and signal transducers, were significantly activated upon loss of KDM2B-CxxC (**Figure 5D-5F**). To validate whether the canonical Wnt signaling was enhanced in cKO hippocampi, *Kdm2b^Emx1-ΔCxxC^* cKO mice were crossed with the BAT-GAL (B6.Cg-Tg(BAT-lacZ)3Picc/J) Wnt-reporter mice. Beta-galactosidase staining showed that P0 cKO hippocampi had stronger canonical Wnt activity, including the CA region and FDJ. The ventricular surface of hippocampi and neocortices of cKO brains also displayed elevated Wnt signaling (**Figure 5G-5I**). In particular, components of canonical Wnt signaling genes such as *Lef1* and *Sfrp2* were elevated. *In situ* hybridization verified that the expression of *Lef1*, the gene encoding the transcriptional co-factor of β-Catenin to activate Wnt signaling, is significantly up-regulated in E16.5 cKO hippocampi (**Figure5J-5L**). LEF1 not only has an early role in specifying the hippocampus, but also controls the generation of dentate gyrus granule cells(Galceran, Miyashita-Lin, Devaney, Rubenstein, & Grosschedl, 2000). Similarly, the expression of *Sfrp2*, another Wnt signaling pathway component was greatly enhanced in E16.5 cKO hippocampi and VZ/SVZ of neocortices (**Figure 5M-5O**). Although members of the SFRP family were first reported as Wnt inhibitors, *Sfrp2* regulates anteroposterior axis elongation, optic nerve development, and cardiovascular and metabolic processes by the promoting or inhibiting Wnt signaling pathway(Guan et al., 2021; Marcos et al., 2015; Satoh, Gotoh, Tsunematsu, Aizawa, & Shimono, 2006; von Marschall & Fisher, 2010),

**Figure 5.**
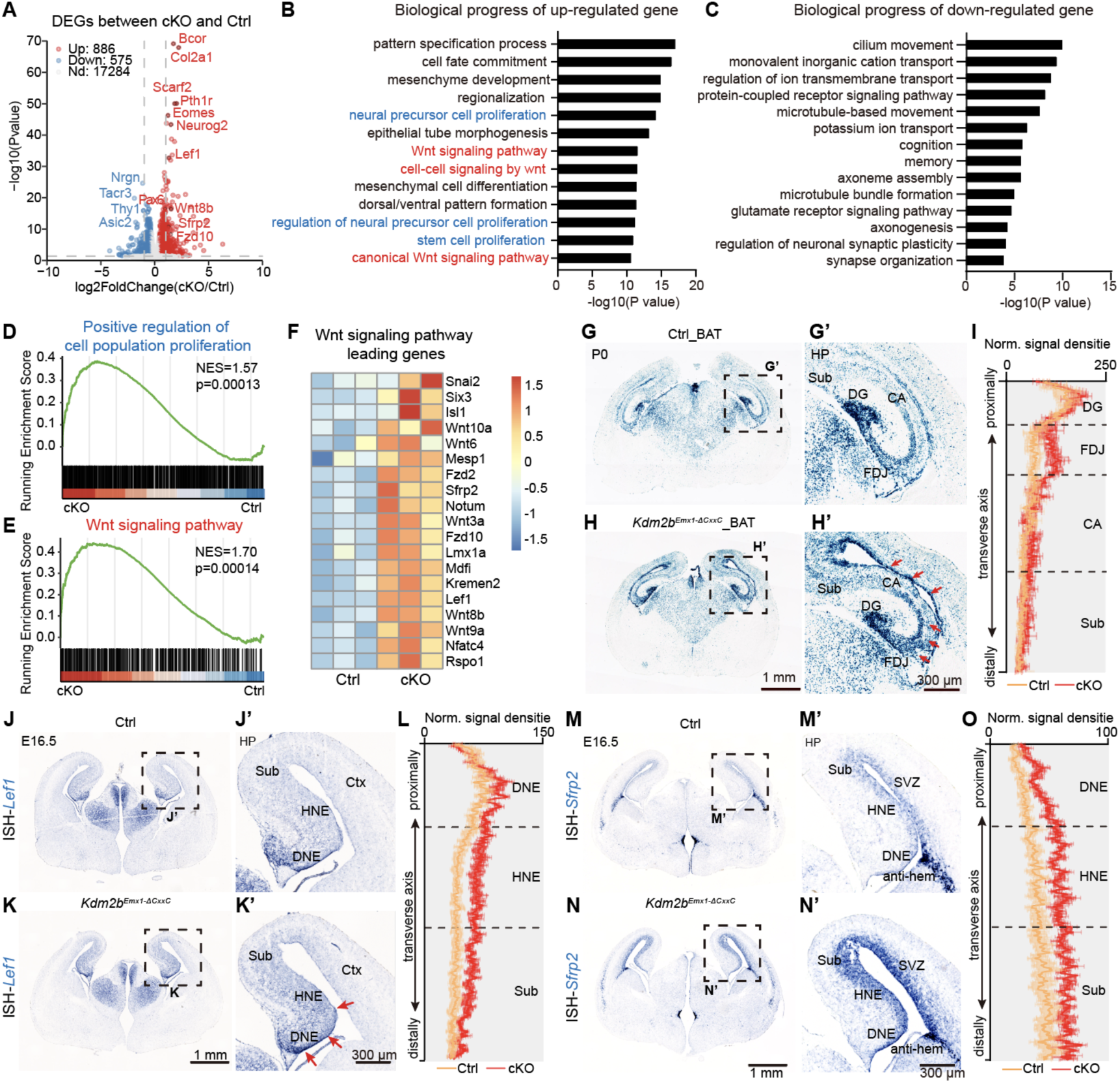
Loss of KDM2B-CxxC results in activation of the Wnt signaling pathway in the hippocampus. (A) The volcano plot of genes up-regulated (red) and down-regulated (blue) in P0 *Kdm2b^Emx1-ΔCxxC^* hippocampi compared to controls. (B) GO analysis of the biological progress of up-regulated genes in P0 *Kdm2b^Emx1-ΔCxxC^* hippocampi revealed terms related to cell proliferation (blue) and Wnt signaling pathways (red). (C) GO analysis of the biological progress of down-regulated genes in P0 *Kdm2b^Emx1-ΔCxxC^* hippocampi. (D) GSEA analysis of positive regulation of cell population proliferation. (E) GSEA analysis of Wnt signaling pathway. (F) The heat map of leading genes in the GSEA of Wnt signaling pathway. (G-H) X-gal staining on P0 coronal sections of Control_BAT (G) and *Kdm2b^Emx1-ΔCxxC^*_BAT (H) brains. Boxed regions were enlarged on the right (G’ and H’). Red arrows indicate areas where X-gal signals are significantly enhanced. (I) Quantification of normalized X-gal signal density on the DG-CA-FDJ-Sub path. n = 4 for Control_BAT brains and n = 3 for *Kdm2b^Emx1-ΔCxxC^*_BAT brains. (J-K) *In situ* hybridization (ISH) of *Lef1* on E16.5 Control (J) and *Kdm2b^Emx1-ΔCxxC^* (K) coronal brain sections, with boxed regions magnified on the right (J’ and K’). Red arrows indicate areas where the *Lef1* expression is significantly elevated. (L) Quantification of normalized ISH signal density of *Lef1* on the DNE-HNE-Sub path. n = 4 for Control_BAT brains and n = 4 for *Kdm2b^Emx1-ΔCxxC^*_BAT brains. (M-N) ISH of *Sfrp2* on E16.5 Control (M) and *Kdm2b^Emx1-ΔCxxC^* (N) coronal brain sections, with boxed regions magnified on the right (M’ and N’). (O) Quantification of normalized ISH signal density of *Sfrp2* on the DNE-HNE-Sub path. n = 4 for Control_BAT brains and n = 4 for *Kdm2b^Emx1-ΔCxxC^*_BAT brains. Scale bars, 1 mm (G, H, J, K, M and N), 300 μm (G’, H’, J’, K’, M’ and N’). HP, Hippocampus; DG, dentate gyrus; FDJ, fimbriodentate junction; CA, Cornu Ammonis; Sub, Subiculum; DNE, dentate neuroepithelium; HNE, hippocampal neuroepithelium; Cx, cortex; SVZ, subependymal ventricular zone.

Since P0 hippocampi contained multiple cell types ranging from RGCs to IPCs to neurons, we propagated RGCs *in vitro* under the serum-free neurosphere condition (**Figure S7A**). RNA-seq transcriptome studies revealed that a significant portion of activated and repressed genes in cKO neurospheres overlapped with those in cKO hippocampal tissue (**Figure S7B-S7E**). GO analyses indicated genes involved in cell division and canonical Wnt signaling were activated in cKO hippocampal neurospheres (**Figure S7F**).

KDM2B is a key component of variant PRC1 to mediate repressive histone modification H2AK119ub and subsequent H3K27me3 (**Figure 6A**) and its long isoform KDM2BLF also bears demethylase activity for H3K36me2. We then examined how loss of KDM2B-CxxC affects these modifications in hippocampal tissue and neurospheres, and whether these effects were associated with enhanced Wnt signaling. To this end, chromatin immunoprecipitation sequencing (ChIP-seq) was performed using P0 hippocampal tissue and neurospheres. As expected, the overall levels of H2AK119ub and H3K27me3 were decreased in cKO hippocampal tissues and neurospheres (**Figure 6B-6C**). However, the level of H3K36me2 was not altered in cKO tissue but slightly increased in neurospheres (**Figure 6B-6C; S8H-S8I**). Moreover, assay for transposase-accessible chromatin using sequencing (ATAC-seq) showed that chromatin became more accessible in cKO hippocampi (**Figure 6C**).

**Figure 6.**
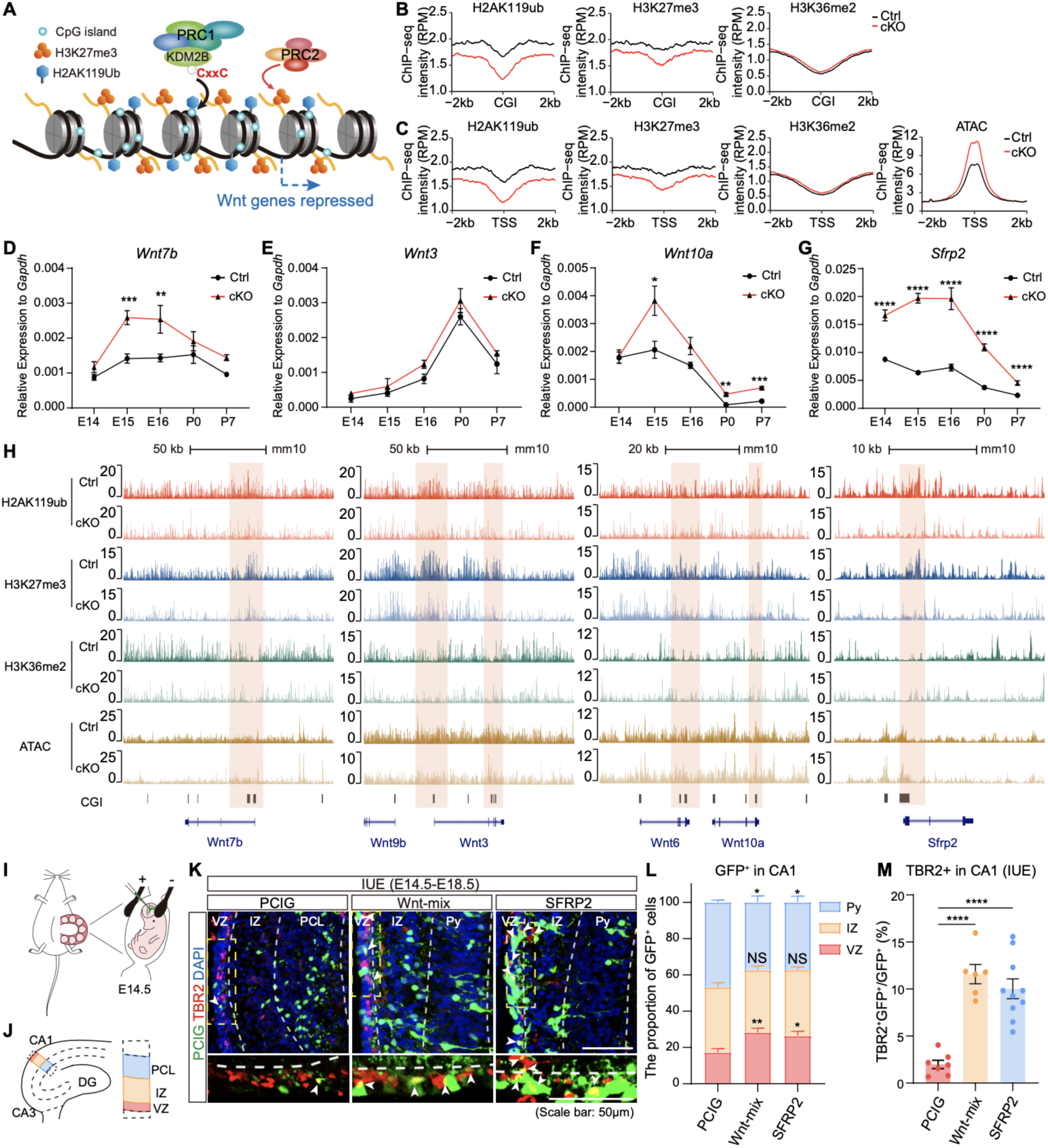
KDM2B epigenetically silences components of Wnt signaling genes in developing hippocampi. (A) The working diagram of KDM2B: KDM2B-CxxC recognizes and binds to CpG islands (CGI) of DNA, therefore recruiting PRC1 to CpG islands (CGIs). Reciprocal recognition of modifications by PRC1 and PRC2 leads to enrichment of H2AK119ub and H3K27me3, hence stabilizing gene repression. (B) Line charts showing average H2AK119ub, H3K27me3 and H3K36me2 signals at CGIs (± 2 kb flanking regions) in P0 control (black lines) and *Kdm2b^Emx1-ΔCxxC^* (cKO) (red lines) hippocampi. (C) Line charts showing average H2AK119ub, H3K27me3, H3K36me2 and ATAC-seq signals at CGI+ TSS (±2 kb flanking regions) in P0 control (black lines) and *Kdm2b^Emx1-ΔCxxC^* (cKO) (red lines) hippocampi. TSS, transcription starting sites. (D-G) RT-qPCR showing relative expressions of *Wnt7b*, *Wnt3*, *Wnt10a* and *Sfrp2* in control (black lines) and *Kdm2b^Emx1-ΔCxxC^* (red lines) hippocampi of indicated developmental stages (E14.5, E15.5, E16.5, P0 and P7). (H) The UCSC genome browser view of HA2K119ub, H3K27me3 and H3K36me2 enrichment and ATAC-seq signal in P0 control and *Kdm2b^Emx1-ΔCxxC^* (cKO) hippocampi at Wnt gene loci [corresponding to (D-G), *Wnt7b*, *Wnt3*, *Wnt10a* and *Sfrp2*]. CGIs were shown as black columns at the bottom, and signals represent ChIP-seq RPM (reads per million). Colored regions marked enrichment differences between control and cKO. (I) The schematic diagram of *in utero* electroporation (IUE) to target the developing hippocampi. (J) The schematic diagram of hippocampal structure, and the hierarchical partition of CA1 region (VZ, IZ, Py). (K) E14.5 mouse hippocampi were electroporated with empty or Wnt-mix-expressing vector (Wnt3a, Wnt5a, Wnt5b, Wnt7b, and Wnt8b) or SFRP2-expressing vector, along with the GFP-expressing vector (PCIG) to label transduced cells. Embryos were sacrificed at E18.5 for immunofluorescent analysis. Representative immunofluorescent images showing expression of TBR2+ (red) in GFP+ (green) transduced cells at E18.5 CA1 regions. Nuclei were labeled with DAPI (blue). Arrowheads denote double-labeled cells. White dashed lines distinguish three layers of CA1: VZ, IZ, Py. The VZ layer marked by the yellow dashed box is enlarged below. (L) The relative location of GFP+ cells in VZ, IZ and Py were quantified. n = 7 for PCIG, n = 6 for Wnt-mix and n = 10 for SFRP2. (M) Quantification of the proportion of TBR2+GFP+/TBR2+ in CA1 of PCIG, Wnt-mix and SFRP2. n = 7 for PCIG, n = 6 for Wnt-mix and n = 10 for SFRP2. Data are represented as means ± SEM. Statistical significance was determined using two-way ANOVA followed by Sidak’s multiple comparisons test (D-G), using two-way ANOVA followed by Tukey’s multiple comparisons test (L), or using one-way ANOVA analysis (M). *P < 0.05, **P < 0.01, ***P < 0.001, and ****P < 0.0001. Scale bars, 50 μm (K). CA, Cornu Ammonis; VZ, ventricular zone; IZ, intermediated zone; Py, pyramidal cell layer of the hippocampus.

We next verified changes of Wnt signaling pathway and investigated how histone modifications and chromatin accessibilities correlate with changes of gene expression. Since PRC1.1 could be recruited to CGIs *via* KDM2B’s CxxC zinc finger to catalyze H2AK119ub, we paid special attention to chromatin status of CGIs. Quantitative reverse transcription PCR (RT-qPCR) revealed that the transcripts of multiple Wnt ligands and *Sfrp2* in cKO hippocampi were significantly elevated throughout developmental time points (**Figure 6D-6G; S8B-S8G**).

Importantly, peaks for H2AK119ub and H3K27me3 were decreased on CGI-enriched promoters of many activated genes, including *Wnt7b*, *Wnt3*, *Wnt6*, *Wnt10a* and *Sfrp2*, as well as *Pax6*, *Eomes* and *Neurod1*; but peaks for H3K36me2 on these sites were not significantly altered in P0 cKO hippocampi (**Figure 6H; S8J**). Consistently, ATAC-seq showed these CGI promoters were more accessible in cKO hippocampi (**Figure 6H; S8J**). For *Lef1*, the enrichment of H2AK119ub and H3K27me3 was diminished around its CGI promoter in cKO neurospheres (**Figure S8K**). To ask whether activating the Wnt pathway could hamper hippocampal neurogenesis, we electroporated a mix of plasmids expressing Wnt ligands into E14.5 hippocampal primordia. Since *Sfrp2* is one of the most enhanced Wnt signaling components upon loss of KDM2B-CxxC, constructs overexpressing *Sfrp2* were separated transduced into E14.5 hippocampal primordia (**Figure 6I-6J**). Data showed that overexpressing Wnt ligands or *Sfrp2* could significantly block hippocampal neurogenesis, as more transduced cells resided in VZ and IZ but fewer in the pyramidal cell layer (Py) compared to controls (**Figure 6K-6L**), with significantly more transduced cell co-expressing TBR2 (**Figure 6M**).

### Loss of Ring1B did not cause accumulation of neural progenitors

KDM2B recruits other component of PRC1.1, including the ubiquitin protein ligase Ring1B, to CGIs to initiate and stabilize gene silencing. We then asked whether the impeded migration and differentiation of neural progenitors in *Kdm2b* cKO hippocampi is caused by PRC1’s loss-of-function. We obtained *Rnf2* (the gene encoding Ring1B) cKO mice - *Rnf2^Emx1-cKO^* – by crossing floxed *Rnf2* mice with *Emx1*-Cre mice (**Figure S9A**). As expected, ablation of *Rnf2* greatly decreased the level of H2AK119ub in P0 neocortical tissues, with levels of H3K27me3 also slightly decreased (**Figure S9B**). We then stained P0 brains with TBR2 (**Figure S9C-S9D**) to find *Rnf2^Emx1-cKO^* hippocampi were smaller than controls and the number of TBR2+ progenitors in *Rnf2^Emx1-cKO^* hippocampi was decreased by 9.2%. Moreover, the distribution of TBR2+ progenitors in DGs, but not DNE and FDJ, was significantly decreased in the *Rnf2^Emx1-cKO^* hippocampi. However, to our surprise, there was no accumulation and dispersion of TBR2+ IPCs in the FDJ region (2ry) of the *Rnf2^Emx1-cKO^* hippocampi as found in *Kdm2b^Emx1-ΔCxxC^* cKO brains (**Figure S9E**). Therefore, although PRC1’s loss-of-function also causes hippocampal agenesis, it did not lead to buildup of neural progenitors in the migrating path of developing hippocampi.

Together, loss of KDM2B-CxxC reduces repressive histone modifications on key Wnt signal genes, hence leading to the block of their attenuation over time, which causes hampered differentiation and migration of hippocampal progenitors (**Graphic abstract**).

## Discussion

The hippocampus is evolutionarily more ancient than the neocortex(Bingman, Salas, & Rodriguez, 2009) and the production and localization of CA pyramidal neurons also follows the birthdate-dependent inside-out pattern(Supèr, Soriano, & Uylings, 1998). In neocortical development, Pax6+ RGCs and TBR2+ IPCs largely reside in the VZ and SVZ respectively, with their nuclei undergoing local oscillation. Although many cellular and epigenetic programs were found to control numbers and differentiation of neocortical RGCs and IPCs, it remains unclear how these mechanisms were applied in hippocampal development, which involves migration and dispersion of neural progenitors. Here we revealed that the chromatin association of KDM2B, an essential component of variant PRC1.1, is required for hippocampal formation. KDM2B mediates silencing of Wnt signaling genes to facilitate proper migration and differentiation of hippocampal progenitors.

Our knowledge on the mammalian Polycomb repressive system has mostly come from studies in pluripotent stem cells and in embryos at early developmental stages(Chen, Djekidel, & Zhang, 2021; Laugesen & Helin, 2014; O’Carroll et al., 2001; Sugishita et al., 2021; Voncken et al., 2003), when the establishment of repressive domains is initiated(Bonnet et al., 2022). Nonetheless, how Polycomb controls sequential fate determination in specific tissues and at later developmental stages largely remains elusive. Both PRC1 and PRC2 are essential players in neocortical development(Eto et al., 2020; Morimoto-Suzki et al., 2014; Pereira et al., 2010; Piper et al., 2014; Sun, Chang, Gerhartl, & Szele, 2018). The deletion of Ring1B, the core enzymatic component of PRC1, prolonged neocortical neurogenesis at the expense of gliogenesis. PRC1 regulates the chromatin status of neurogenic genes of neural progenitors, hence altering their responsiveness to neurogenic Wnt signals over developmental time(Hirabayashi et al., 2009). Ring1B was also found to regulate dorsoventral patterning of the forebrain(Eto et al., 2020) and sequential production of deep and upper-layer neocortical PNs(Morimoto-Suzki et al., 2014). However, we surprisingly revealed that ablation of Ring1B did not significantly hamper migration and distribution of TBR2+ neural progenitors of developing hippocampi, whereas these defects are prominent in the *Kdm2b^Emx1-ΔCxxC^* hippocampi. In addition, removal of KDM2B from the chromatin only have mild effects on neocortical development and overall H2AK119ub1 and H3K27me3 levels. Therefore, KDM2B likely controls fate determination of hippocampal progenitors by selectively repressing a series of progenitor genes including those in the Wnt pathway *via* PRC1.1.

Knocking out EED, one of the core components of PRC2, results in hampered neurogenesis of postnatal DG. However, unlike *Kdm2b* cKO brains, the EED knockouts did not display any hippocampal malformation at P0(Liu et al., 2019). Moreover, deletion of *Ezh2* in adult NSCs leads to disturbed neurogenesis of DG(Zhang et al., 2014), which was unseen in *Kdm2b* cKO DGs. These discrepancies echoes either distinct roles or spatiotemporal activities of PRC1 and PRC2. It would be worthy of dissecting functions of distinct PRC1/2 variants or their components in neural development and homeostasis(Lan et al., 2022). A number of studies indicated that PRC1 and PRC2 can directly or indirectly regulate Wnt signaling in developmental, physiological and disease conditions(Chiacchiera et al., 2016; Oittinen et al., 2016; Jiajia Wang et al., 2020). Wnt signaling governs multiple aspects of neural development including neurulation, pattern formation, and fate choices of neural progenitors(Bengoa-Vergniory & Kypta, 2015; Chenn & Walsh, 2002; Machon, van den Bout, Backman, Kemler, & Krauss, 2003). Moreover, the strength and gradient of the canonical Wnt signaling in the RGC-IPC-neuron path and through developmental time ensures proper cell fate establishment and transition(Lie et al., 2005; Munji, Choe, Li, Siegenthaler, & Pleasure, 2011; Junbao Wang et al., 2022), including those in hippocampal morphogenesis(Galceran et al., 2000; Zhong et al., 2020). Deletion of KDM2B-CxxC greatly elevated Wnt signaling in hippocampi of multiple developing stages and in hippocampal progenitors, which could lead to impeded migration and differentiation of IPCs. Of note, however, the ablation of KDM2B-CxxC has minimal effects on neocortical development, reflecting regional difference between neocortical and hippocampal progenitors.

The deletion of KDM2B-CxxC would totally abolish KDM2B’s association with CGIs, thus disabling KDM2B’s two major functions on chromatin - mediating H2AK119Ub *via* PRC1.1 and demethylating H3K36me2, the latter of which is solely executed by the JmjC-containing KDM2BLF. Previous studies indicated that the demethylase activity of KDM2A/B is required for PRC establishment at CGIs of peri-implantation embryos(Huo et al., 2022), but contributes moderately to the H3K36me2 state at CGI-associated promoters and is dispensable for normal gene expression in mouse embryonic stem cells(Turberfield et al., 2019). Interestingly, levels of H3K36me2 were not significantly altered globally or locally in KDM2B-CxxC deleted hippocampi. Moreover, *Kdm2blf^KO^* mice did not display hippocampal agenesis or malformation(Fukuda, Tokunaga, Sakamoto, & Yoshida, 2011; W. Li et al., 2020). Thus, the H3K36me2 demethylase activity of KDM2B is likely dispensable for hippocampal development.

*KDM2B* is implicated in neurological disorders including ID and behavior abnormalities, and the region encoding the CxxC ZF is the mutational hotspot(van Jaarsveld et al., 2022). Consistently, *Kdm2b^Emx1-ΔCxxC^* cKO mice displayed prominent defects in spatial and motor learning and memory, as well as contextual fear conditioning. It would be essential to explore whether patients with *KDM2B* mutations have defects of hippocampal morphogenesis and function, and how KDM2B mediated gene repression is implicated in human brain development.

## MATERIALS AND METHODS

### Mice and genotyping

All animal procedures were approved by the Animal Care and Ethical Committee of Wuhan University. Wild-type CD-1 (ICR) and C57BL/6 mice were obtained from the Hunan SJA Laboratory Animal Company (Changsha, China). Mice were housed in a certified specific-pathogen-free (SPF) facility. The noon of the day when the vaginal plug was found was counted as embryo 1. (E) day 0.5.

Mice with conditional deletion of *Kdm2b-CxxC* were obtained by first crossing *Kdm2b^fl/fl^* (generated by Applied Stem Cell) females with *Emx1-Cre* (Jackson Laboratories, stock number 005628), *Nestin-Cre* (Jackson Laboratories, stock number 003771) or *Nex-Cre* males [*Neurod6^tm1(cre)Kan^*, MGI:2668659]. *Emx1-Cre; Kdm2b^fl/+^*, *Nestin-Cre; Kdm2b^fl/+^* or *Nex-Cre; Kdm2b^fl/+^* males were crossed with *Kdm2b^flfl^* females to obtain conditional knockout mice (*Kdm2b^Emx1-ΔCxxC^*, *Kdm2b^Nestin-ΔCxxC^*, *Kdm2b^Nex-ΔCxxC^*). *Kdm2b^fl/+^* and *Kdm2b^fl/fl^* were phenotypically indistinguishable from each other, and used as controls. The primer set forward 5’-cctgtagtccttggtatttcctggc-3’/reverse 5’ - cccaacttgcccttaggccg-3’ was used for mice genotyping, and band sizes for *Kdm2b^fl/+^* mice are 364 bp (WT allele) and 404 bp (targeted allele with 5’ loxP). Forward 5’-cctgttacgtatagccgaaa-3’/reverse 5’-cttagcgccgtaaatcaatc-3’ was used for *Emx1-Cre*, *Nestin-Cre* and *Nex-Cre* genotyping with band size 319 bp (Cre allele).

To analyse adult neurogenesis, the *Nestin-CreERT2* (Jackson Laboratories, stock number 016261) and *Kdm2b^fl/fl^* mice were crossed to generate *Nestin-CreERT2; Kdm2b^fl/+^* animals. *Nestin-CreERT2; Kdm2b^fl/+^* mice were further crossed with *Kdm2b^fl/fl^* mice to obtain homozygous *Kdm2b^NestinCreERT2-ΔCxxC^* animals, which were used for the experiment. Forward 5’-gaccaggttcgttcactca-3’/reverse 5’-caagttaggagcaaacagtagc-3’ was used for *Nestin-CreERT2* genotyping with band size 993 bp (CreERT2 allele).

BAT-Gal mice were kind gifts from Dr. Junlei Chang (Jackson Lab, stock number 005317). To explore the Wnt signaling pathway in *Kdm2b^Emx1-ΔCxxC^* mice, the BAT-Gal and *Kdm2b^fl/fl^* mice were crossed to generate *Kdm2b^fl/+^*; BAT-Gal animals. *Emx1-Cre; Kdm2b^fl/+^* mice were further crossed with *Kdm2b^fl/+^*; BAT-Gal mice to obtain *Kdm2b^Emx1-ΔCxxC^*; BAT-Gal animals, which were used for the experiment. The primer set forward 5′-atcctctgcatggtcaggtc-3′/reverse 5′-cgtggcctgattcattcc-3′ was used for BAT-Gal mice with band size 315 bp (LacZ allele).

Mice with conditional deletion of *Rnf2* were obtained by first crossing *Rnf2^fl/fl^* (purchased from GemPharmatech, Strain NO. T014803) females with *Emx1-Cre* males (Jackson Laboratories, stock number 005628). The primer set forward 5’-agctgtggtcctgcgtttcatttc-3’/reverse 5’ - gctcttactgtgttacaaccctagccc-3’ was used for *Rnf2^fl/+^* mice genotyping, and band sizes for *Rnf2^fl/+^* mice are 289 bp (WT allele) and 391 bp (targeted allele with 5’ loxP).

### Tamoxifen and BrdU administration

To activate Cre-mediated recombination, tamoxifen (TAM; Sigma-Aldrich) was used, which was made fresh daily and dissolved in sunflower oil solution (Sigma-Aldrich). 8-week-old *Kdm2b^NestinCreERT2-ΔCxxC^* mice were daily administered with 30 mg/kg prewarmed TAM intraperitoneally for 6 consecutive days (d1-d6). From day 2 to day 7, mice were injected with 50 mg/kg BrdU (Sigma-Aldrich) intraperitoneally for 6 consecutive days and were sacrificed 1 day later (day 8, Short-Term) or 4 weeks later (day 35, Long-Term) to identify BrdU-positive adult-born cells.

### Tissue fixation and sectioning

The pregnant dam was anesthetized with 0.7% w/v pentobarbital sodium (105 mg/kg body weight) in 0.9% sodium chloride. Embryos were sequentially removed from the uterus. Brains of embryos were dissected out in cold PBS and immersed in 4% paraformaldehyde (PFA) overnight at 4°C. For P0, P2, P7 and adult mice, animals were anesthetized with 0.7% w/v pentobarbital sodium solution followed by trans-cardiac perfusion with 4% PFA in PBS (P0, 5 ml; P7, 10 ml; adult, 30 ml). Brains were dissected and post-fixed in 4% PFA overnight at 4°C. Next day, brains were dehydrated in 20% w/v sucrose overnight at 4°C. For sectioning, brains were embedded in OCT (SAKURA) and cut at 20 μm for adult brains and 14 μm for other stages with a cryostat (Leica CM1950).

### Nissl staining

Adult brain sectionswere stained with 0.25% Cresyl Violet (Sigma-Aldrich) solution for 15 min at 65°C. Sections were then decolorized in ethanol for 0.5-1 min, dehydrated in gradient ethanol solutions for 5 min each and cleared twice in xylene for 5 min. Sections were mounted in the neutral balsam.

#### *In situ* hybridization (ISH)

Sections were dried in a hybridization oven at 50°C for 15 min and fixed in 4% PFA for 20 min at room temperature, followed by permeabilization in 2 μg/ml proteinase K in PBS for 10 min at room temperature. Prior to hybridization, sections were acylated in 0.25% acetic anhydride for 10 min. Then, sections were incubated with a digoxigenin-labeled probe diluted (0.2 ng/μl) in hybridization buffer (50% deionized formamide, 5× SSC, 5× Denhart’s, 250 μg/ml tRNA, and 500 μg/ml Herring sperm DNA) under coverslips in a hybridization oven overnight at 65°C. The next day, sections were washed 4 times for 80 min in 0.1× SSC at 65°C. Subsequently, they were treated with 20 μg/ml RNase A for 20 min at 37°C, then blocked for 3.5 h at room temperature in 10% normal sheep serum. Slides were incubated with 1:5,000 dilution of anti-digoxigenin-AP conjugated antibody (Roche) overnight at 4°C. BCIP/NBT (Roche) was used as a color developing agent. ISH primers used are listed in Table S1.

### Immunofluorescence

Frozen brain sections were mounted onto Superfrost plus slides and then dried at room temperature. For heat-mediated antigen retrieval, slides were incubated for 15 min in 10 mM sodium citrate buffer (pH 6.0) at 95°C. For BrdU staining, sections were treated with 20 μg/ml proteinase K (Sigma) (1:1000 in PBC) for 5 min and 2 N HCl for 30 min at room temperature. Sections were then immersed in blocking buffer (3% normal sheep serum and 0.1% Triton X-100 in PBS; or 5% BSA and 0.5% Triton X-100 in PBS) for 2 h at room temperature. Sections were then incubated in primary antibodies [mouse anti-Calbindin (1:1000; Sigma, C9848), rabbit anti-ZBTB20 (1:1000; Sigma, HPA016815), mouse anti-HopX (1:200; Santa Cruz, sc-398703), rabbit anti-Wfs1 (1:1000; Proteintech, 86995), mouse anti-PROX1 (1:200; Millipore, MAB5654), rabbit anti-GFAP (1:500; DAKO, Z0334), rat anti-CTIP2 (1:500; Abcam, ab18465), rabbit anti-SATB2 (1:500; Abcam, ab92446), rat anti-BrdU (1:500; Abcam, ab6326), mouse-anti-BrdU (1:500; Roche, 11170376001), rabbit-anti-DCX (1:500; Abcam, ab18723), rabbit anti-TBR2 (1:500; Abcam, ab23345), rat anti-TBR2 (1:500; Thermo Fisher, 14-4875-82), rabbit anti-PAX6 (1:500; Millipore, ab2237), and rabbit anti-Ki67 (1:500; Abcam, ab15580] in blocking buffer overnight at 4°C. After three rinses in PBS, sections were incubated in secondary antibodies (Alexa Fluor 488-conjugated anti-mouse, A11029; Alexa Fluor 555-conjugated anti-mouse, A21422; Alexa Fluor 488-conjugated anti-rat, A11006; Alexa Fluor 555-conjugated anti-rat, A21434; Alexa Fluor 647-conjugated anti-rat, A21247; Alexa Fluor 488-conjugated anti-rabbit, A11034; Alexa Fluor 555-conjugated anti-rabbit, A21429; Alexa Fluor 647-conjugated anti-rabbit, A21245; Alexa Fluor 488-conjugated anti-chicken, A11039; Thermo Fisher Scientific; 1:1000) for 1 h at room temperature. Nuclei were labeled by incubation in PBS containing 4′,6-diamidino-2-phenylindole (DAPI) (0.1 μg/ml) (Sigma-Aldrich), and samples were mounted in ProLong Gold Antifade Mountant (Thermo Fisher Scientific).

### 5-Ethynyl-2′-Deoxyuridine (EdU) staining

Proliferation of cells was investigated with BeyoClickTM EdU Cell Proliferation Kit (C0075S, Beyotime, China) according to the manufacturer’s protocols. In brief, frozen brain sections were dried at temperature and permeated with 0.3% Triton X-100 in PBS for 30 min. Sections were then incubated with EdU working solution for 1 h at 37 °C in the dark. After incubation, regular immunofluorescence staining can be followed.

### Immunohistochemical staining

Frozen brain sections were dried at room temperature, and then pretreated with 0.3% H_2_O_2_ for 15 min to deactivate endogenous peroxidase. Sections were blocked with 3% normal sheep serum with 0.1% Tween 20 at room temperature for 2 h. Sections were then incubated in primary antibodies [rabbit anti-NeuN (1:500; Abcam, ab177487), rabbit-anti-DCX (1:500; Abcam, ab18723), rabbit anti-SOX2 (1:500; Millipore, ab5603), rabbit anti-TBR2 (1:500; Abcam, ab23345), and rabbit anti-BLBP (1:500; Abcam, ab32423)] in blocking buffer overnight at 4°C, followed by addition of the avidin-biotin-peroxidase complex (1:50; VECTASTAIN Elite ABC system, Vector Laboratories). Peroxidase was reacted in 3,3′-diaminobenzidine (5 mg/ml) and 0.075% H_2_O_2_ in Tris-HCl (pH 7.2). Sections were dehydrated in gradient ethanol (75% ethanol, 95% ethanol, 100% ethanol and 100% ethanol, each for 5 min), and cleared twice in xylene for 5 min, then mounted in the neutral balsam.

### Behavior tests

We used 12-to 16-week-old age-matched male mice for all behavioral tests. Mice were housed (3-5 animals per cage) in standard filter-top cages with access to water and rodent chow at all times, maintained on a 12:12 h light/dark cycle (09:00-21:00 h lighting) at 22°C, with relative humidity of 50%–60%. All behavioral assays were done blind to genotypes.

*Open field test.* The test mouse was gently placed near the wall-side of a length of 50 cm, a width of 50 cm, and a height of 50 cm open-field arena and allowed to explore freely for 20 min. Only the last 10 min of the movement of the mouse was recorded by a video camera and analyzed with Ethovision XT 13 (Noldus).

*Rotarod test.* The test consists of 4 trials per day for 10 days with the rotarod (3 cm in diameter) set to accelerate from 4 rpm to 40 rpm over 5 minutes. The trial started once mice were placed on the rotarod rotating at 4 rpm in partitioned compartments. The time for each mouse spent on the rotarod were recorded. At least 20 min recovery time was allowed between trials. The rotarod apparatus was cleaned with 70% ethanol and wiped with paper towels between each trial.

*Morris water maze.* Mice were introduced into a stainless water-filled circular tank, which is 122 cm in diameter and 51 cm in height with non-reflective interior surfaces and ample visual cues. Two principal axes were drawn on the floor of the tank, each line bisecting the maze perpendicular to one another to create an imaginary ‘+’. The end of each line demarcates four cardinal points: North, South, East and West. To enhance the signal-to-noise ratio, the tank was filled with water colored with powdered milk. A 10-cm circular plexiglass platform was submerged 1 cm below the surface of the water in middle of the southwest quadrant. Mice started the task at fixed points, varied by day of testing(Vorhees & Williams, 2006). Four trials were performed per mouse per day with 20 min intervals for 5 days. Each trial lasted 1 min and ended when the mouse climbed onto and remained on the hidden platform for 10 s. The mouse was given 20 s rest on the platform during the inter-trial interval. The time taken by the mouse to reach the platform was recorded as its latency. Times for four trials were averaged and recorded as a result for each mouse. On day 6, the mouse was subjected to a single 60-s probe trial without a platform to test memory retention. The mouse started the trial from northeast, the number of platform crossings was counted, and the swimming path was recorded and analyzed using the Ethovision XT 13 (Noldus).

*Fear conditioning (FC).* The FC apparatus consisted of a conditioning box (18 × 18 × 30 cm), with a grid floor wired to a shock generator surrounded by an acoustic chamber and controlled by Ethovision XT 13 (Noldus). On Day 1, each mouse was placed in the conditioning box for 2 min, and then a pure tone (80 db) was sounded for 30 s followed by a 2 s foot shock (0.4 mA). Two minutes later, this procedure was repeated. After the delivery of the second shock, mice were returned to their home cages. On Day 2, each mouse was first placed in the fear conditioning chamber containing the exact same context, but there was no administration of a tone or foot shock. Freezing was analyzed for 4 min. One hour later, the mice were placed in a new context (containing a different odor, cleaning solution, floor texture, walls and shape) where they were allowed to explore for 3 min before being re-exposed to the fear conditioning tone and freezing was assessed for an additional 3 min. FC was assessed through the continuous measurement of freezing (complete immobility), which is the dominant behavioral fear response. Freezing was measured using the Noldus Ethovision video tracking system (Ethovision XT 13).

*Forced swimming test.* For the forced swimming test, the test mouse was placed into a 20 cm height and 17 cm diameter glass cylinder filled with water to a depth of 10 cm at 22°C. The test continues 6 min and the immobility time of the last 5 minutes was recorded for further processing.

*Tail suspension test.* The test mouse was suspended in the middle of a tail suspension box (55 cm height × 60 cm width × 11.5 cm depth) above the ground by its tail. The mouse tail was adhered securely to the suspension bar using adhesive tapes. After 1 min accommodation, the immobility time was recorded by a video camera and analyzed by Ethovision XT 13 (Noldus).

*Elevated plus maze test.* The elevated plus maze, made of gray polypropylene and elevated about 40 cm above the ground, consists of two open arms and two closed arms (each 9.5 cm wide and 40 cm long). To assess anxiety, the test mouse was placed in the central square facing an open arm and allowed to explore freely for 5 min. The time spent in the open arm was analyzed with the Ethovision XT 13 (Noldus).

### X-Gal staining

Frozen sections were fixed in fresh cold fixative (0.2% PFA) in buffer L0 (0.1M PIPES buffer (pH 6.9), 2mM MgCl_2_, 5mM EGTA) for 10 min. Slides were rinsed in PBS plus 2mM MgCl_2_ on ice, followed by a 10 min wash in the same solution. Place slides in detergent rinse [0.1M PBS (pH 7.3), 2mM MgCl_2_, 0.01% sodium deoxycholate, 0.02% Nonidet P-40] on ice for 10 min. Slides were then moved to a freshly made and filtered X-Gal staining solution [0.1M PBS (pH 7.3), 2mM MgCl_2_, 0.01% sodium deoxycholate, 0.02% Nonidet P-40, 5mM K_3_Fe(CN)_6_, 5mM K_4_Fe(CN)_6_·3H_2_O and 1 mg/ml X-Gal]. Sections were incubated at 37 °C from a few minutes to overnight in the dark. Sections were rinsed with water to stop the reaction. Sections were dehydrated with gradient ethanol and xylene sequentially, and mounted with the neutral balsam.

### RNA isolation and reverse transcription (RT)

RNA isolation was performed using the RNAiso Plus (TAKARA) according to manufacturer’s instructions. Tissue or cells were homogenized using a glass-Teflon in 1 ml or 500 µl RNAiso Plus reagent on ice and phase separation was achieved with 200 µl or 100 µl chloroform. After centrifugation at 12,000× g for 15 min at 4°C, RNA was precipitated by mixing aqueous phase with equal volumes of isopropyl alcohol and 0.5 µl 20 mg/ml glycogen. Precipitations were dissolved in DNase/RNase-free water (not diethylpyrocarbonate treated, Ambion). 1 µg of total RNA was converted to cDNA using M-MLV reverse transcriptase (TAKARA) under standard conditions with oligo(dT) or random hexamer primers and Recombinant RNase Inhibitor (RRI, TAKARA). Then the cDNA was subjected to quantitative RT-PCR (qRT-PCR) using the SYBR green assay with 2× SYBR green qPCR master mix (Bimake). Thermal profile was 95 °C for 5 min and 40 cycles of 95 °C for 15 sec and 60 °C for 20 sec. *Gapdh* was used as endogenous control genes. Relative expression level for target genes was normalized by the Ct value of *Gapdh* using a 2^−ΔΔCt^ relative quantification method. Reactions were run on a CFX Connect TM Real-Time PCR Detection System (Bio-Rad). The primers used are listed in Table S2.

### Neurosphere culture

Mouse hippocampal neural progenitor cells (NPCs) were enriched from P0 mouse hippocampi, cultured on ultra-low-attachment plates (Corning, New York, United States) and maintained in indicated culture media (DMEM/F12, Life Technologies) containing N2 and B27 supplements (1×, Life Technologies), 1mM Na-pyruvate, 1 mM N-acetyl-L-cysteine (NAC), human recombinant FGF2, and EGF (20 ng/mL each; Life Technologies). After cultured *in vitro* for three generations, neurospheres were subjected to RNA-seq and ChIP-seq analyses.

### RNA-seq library construction

Total RNA was extracted as described above. The concentration and quality of RNA was measured with Nanodrop 2000c (Thermo Fisher Scientific) and an Agilent 2100 Bioanalyzer (Agilent Technologies), respectively. RNA-seq libraries were constructed by NEBNext® Ultra™ II RNA Library Prep Kit for Illumina® (NEB #E7775). Briefly, mRNA was extracted by poly-A selected with magnetic beads with poly-T and transformed into cDNA by first and second strand synthesis. Newly synthesized cDNA was purified by AMPure XP beads (1:1) and eluted in 50 μl nucleotide-free water. RNA-seq libraries were sequenced by Illumina NovaSeq 6000 platform with pair-end reads of 150 bp. The sequencing depth was 60 million reads per library.

### Bulk RNA-Seq data analysis

P0 hippocampus and neurosphere RNA-seq data were checked for quality control by FastQC (version 0.11.9). Paired-end reads were trimmed to remove adaptors and low-quality reads and bases using cutadapt (version 3.2). Clean reads were aligned to the mouse UCSC mm10 genome using STAR (version 2.7.10b) with default parameters. The number of covering reads were counted using featureCounts (version v2.0.1). The resulting read counts were processed with R package DESeq2 (version 1.38.1) to identify differential expression genes (log2 fold change > 0.4 and p value < 0.05) between datasets. Cufflinks package (version 2.2.1) assembles individual transcripts from reads that have been aligned to reference genome. The gene expression level was normalized by fragments per kilobase of bin per million mapped reads (FPKM). Gene Ontology (GO) analysis and Gene Set Enrichment Analysis (GSEA) in this study were performed using clusterProfiler (version 4.2.2).

### Chromatin immunoprecipitation (ChIP) assay

For each experiment, single-cell suspensions from P0 hippocampi were collected as described above. The hippocampal tissue was digested into single cells by Papain (20U dissolved in each mL of DMEM/F12 medium, preheated at 37°). Cells were cross-linked with 1% formaldehyde for 10 minutes at room temperature, and quenched with 0.125 M of glycine for 5 minutes. Cross-linked samples were then rinsed twice in PBS, then cells were collected by centrifugation. Next, cells were pretreated with lysis buffer (50 mM of Tris-HCl [pH 8.0], 0.1% SDS, and 5 mM of EDTA) and incubated for 5 minutes with gentle rotation at 4°C. After centrifugation, bottom cells were washed for two times with ice-cold PBS and harvested in ChIP digestion buffer (50 mM of Tris-HCl [pH 8.0], 1 mM of CaCl_2_, and 0.2% Triton X-100). DNA was digested to 150-300 bp by micrococcal nuclease (NEB; M0247S). Sonicate cells in EP tubes with power output 100 W, 5 min, 0.5 s on, 0.5 s off on ice. The resulting lysate was centrifugation and divided into four parts for 10% input, H2AK119Ub, H3K27me3, and H3K36me2 immunoprecipitation. After diluting each sample (in addition to input) to 1 mL with dilution buffer (20 mM of Tris-HCl [pH 8.0], 150 mM of NaCl, 2 mM of EDTA, 1% Triton X-100, and 0.1% SDS), immunoprecipitation was further performed with sheared chromatin and 3 μg rabbit anti-H2AK119Ub antibody (CST, 8240S); or rabbit anti-H3K27me3 antibody (CST, 9733S); or rabbit anti-H3K36me2 antibody (CST, 2901S), then incubated with protein A/G beads overnight at 4°C on a rotating wheel. The next day, beads were wash with Wash Buffer I (20mM Tris-HCl, pH 8.0; 1% Triton X-100; 2mM EDTA; 150mM NaCl; 0.1% SDS), Wash Buffer II (20mM Tris-HCl, pH 8.0; 1% Triton X-100; 2mM EDTA; 500mM NaCl; 0.1% SDS), Wash Buffer III (10mM Tris-HCl, pH 8.0; 1mM EDTA; 0.25M LiCl; 1% NP-40; 1% deoxycholate) and TE buffer. DNA was eluted by ChIP elution buffer (0.1 M of NaHCO_3_, 1% SDS, 20 µg/mL of proteinase K). The elution was incubated at 65°C overnight, and DNA was extracted with a DNA purification kit (DP214-03; TIANGEN).

### ChIP-seq library construction

ChIP-seq libraries were constructed by VAHTS Universal DNA Library Prep Kit for Illumina V3 (Vazyme ND607). Briefly, 50 μL purified ChIP DNA (5 ng) was end-repaired for dA tailing, followed by adaptor ligation. Each adaptor was marked with a barcode of 6 bp which can be recognized after mixing different samples together. Adaptor-ligated ChIP DNA was purified by VAHTS DNA Clean Beads (Vazyme N411) and then amplified by PCR of 10 cycles with primers matching with adaptors’ universal part. Amplified ChIP DNA was purified again using VAHTS DNA Clean Beads in 35-μL EB elution buffer. For multiplexing, libraries with different barcode were mixed together with equal molar quantities by considering appropriate sequencing depth (about 30 million reads per library). Libraries were sequenced by Illumina Nova-seq 6000 platform with pair-end reads of 150 bp.

### ChIP-seq data analysis

DNA libraries were sequenced on Illumina NovaSeq 6000 platform. All P0 hippocampus and neurosphere ChIP-seq data were checked and removed with adaptor sequences same as the RNA-seq data processing. Clean reads were aligned to the mouse UCSC mm10 genome using Bowtie2 (version 2.4.5). Duplicates were removed using the samtools rmdup module. Regions of peaks were called using the SICER software package, with the input genomic DNA as a background control (parameters: -w 200 -rt 1 -f 150 -egf 0.77 -fdr 0.01 -g 600 -e 1000 --significant_reads). The bigwig signal files were visualized using the computeMatrix, plotHeatmap, plotProfile modules in Deeptools (version 3.5.1). Homer was used to identify adjacent genes from the peaks obtained from SICER.

### ATAC-seq library construction

ATAC-seq libraries were constructed by TruePrep DNA Library Prep Kit V2 for Illumina (Vazyme TD501). Briefly, P0 hippocampi were dissected and gently homogenized in cold nuclear isolation buffer (10 mM Tris-HCl, pH 7.4, 10 mM NaCl, 3 mM MgCl_2_, 0.1% Igepal CA-630). Nuclei were collected by centrifugal precipitation. 50,000 nuclei were put into the tagmentation reaction for each sample (performed with 30 min incubation time at 37°C). Immediately following the tagmentation, DNA fragments were purified using VAHTS DNA Clean Beads (2X). Purified DNA fragments were added with Illumina i5+i7 adapters with unique index to individual samples followed by PCR reaction (PCR program: 72°C for 3 min, 98°C for 30 s, 98°C for 15 s, 60°C for 30 s, 72°C for 30 s, repeat 3-5 for 13 cycles, 72°C for 5 min, and hold at 4°C). Generated libraries were purified using VAHTS DNA Clean Beads (1.2X). For multiplexing, libraries with different barcode were mixed together with equal molar quantities by considering appropriate sequencing depth (about 50 million reads per library). Libraries were sequenced by Illumina Nova-seq 6000 platform with pair-end reads of 150 bp.

### ATAC-seq data analysis

P0 hippocampus ATAC-seq raw data were trimmed by Cutadapt with parameters -u 3 -u -75 -U 3 -U -75 -m 30 and then aligned to mouse mm10 genome using Bowtie2 (-X 2000 --very-sensitive). Subsequently, we downloaded blacklisted regions including a large number of repeat elements in the genome from ENCODE project and then removed these significant background noise. ATAC-seq datasets contained a large percentage of reads that were derived from mitochondrial DNA. We removed mitochondrial reads after alignment using Samtools. Then, we filtered reads to remove exact copies of DNA fragments that arise during PCR using Picard’s MarkDuplicates (version 2.26.4). All reads aligning to the + strand were offset by +4 bp, and all reads aligning to the – strand were offset −5 bp, since Tn5 transposase has been shown to bind as a dimer and insert two adaptors separated by 9 bp. We adjusted the shift read alignment using alignmentSieve. Next, peaks calling was finished by Macs2 (version 2.2.7.1) with parameters -f BAMPE --nomodel -- keep-dup all --shift -100 --extsize 200 -g mm --cutoff-analysis -B. We created bigwig files for visualizing using bamCoverage (parameters: --normalizeUsing RPGC –effectiveGenomeSize 2407883318) in deepTools. Homer took narrow peak files as input and checked for the enrichment of both known sequence motifs and de novo motifs.

### Defining genomic features

Mm10 CpG island (CGI) regions were downloaded from the UCSC genome browser database. Promoters were defined as all mouse UCSC mm10 gene TSSs, extended by 2 kb upstream and downstream. CGIs in promoters were defined as ± 2 kb around CGI centers overlap with promoter regions. Overlapped regions between CGIs and promoters were identified using bedtools (version 2.29.2) intersect with parameters -e -f 0.5 -F 0.5. H2AK119ub1 +/-promoter genomic loci were defined as such promoter location with or without H2AK119ub1.

### Data and code availability

The GEO accession number for the RNA-seq, ChIP-seq and ATAC-seq data reported in this paper is GSE222465. RNA-seq data of P0 hippocampus and P0 neurosphere have been deposited at GEO: GSE222464. ChIP-seq and ATAC-seq data of P0 hippocampus and P0 neurosphere in this study have been deposited at GEO: GSE222463 and GSE222462. All data are publicly available as of the date of publication. Custom codes were described in detail at methods part. Any additional information required to analyze the data in this paper is available from authors upon reasonable request.

#### *In utero* electroporation (IUE)

*In utero* microinjection and electroporation were performed as followed. Pregnant CD-1 mice with E14.5 embryos were anesthetized by injection of pentobarbital sodium (70 mg/kg), and the uteri were exposed through a 2 cm midline abdominal incision. Embryos were carefully pulled out using ring forceps through the incision and placed on sterile gauze wet with 0.9% sodium chloride. Plasmid DNA (prepared using Endo Free plasmid purification kit, Tiangen) mixed with 0.05% Fast Green (Sigma) was injected through the uterine wall into the telencephalic vesicle using pulled borosilicate needles (WPI). For gain-of-function experiments, pCIG (1 μg/μl) was mixed with pCAGGS-Wnt3a, pCAGGS-Wnt5a, pCAGGS-Wnt5b, pCAGGS-Wnt7b, pCAGGS-Wnt8b, (Wnt-mix) (0.5 μg/μl each), or with pCAGGS-SFRP2 (2 μg/μl). Control mice were injected with pCIG (1 μg/μl). Five electric pulses (33 V, 50 ms duration at 1 s intervals) were generated using CUY21VIVO-SQ (BEX) and delivered across the head of embryos using 5 mm forceps-like electrodes (BEX). The uteri were then carefully put back into the abdominal cavity, and both peritoneum and abdominal skin were sewed with surgical sutures. The whole procedure was completed within 30 min. Mice were warmed on a heating pad until they regained consciousness and were treated with analgesia (ibuprofen in drinking water) until sacrifice at E18.5.

### Plasmid construction

Full-length mouse *Wnt3a* and *Wnt8b* were amplified from cDNAs of E14.5 mouse hippocampi and then cloned into pCAGGS. Full-length *Wnt5a*, *Wnt5b*, *Wnt7b* were amplified from cDNAs of E16.5 mouse cortex, and then cloned into pCAGGS. Full-length mouse *Sfrp2* was amplified from cDNAs of P0 mouse cortex and then cloned into pCAGGS. The primers used are listed in Table S3.

### Quantification and statistical analysis

Sections used for quantification were position-matched for control and experimental brains. Images were binned against proximal-distal transverse axis to quantify the intensity of ISH or LacZ signals of hippocampi. A plot of normalized average signal intensity with standard error of the mean across those regions was generated using ImageJ.

Statistical tests were performed using GraphPad Prism (version 8.0.2). Data analyzed by unpaired two-tailed t-test were pre-tested for equal variance by F-tests. Unpaired Student’s t-tests (two-tailed) were chosen when the data distributed with equal variance. For normally distributed data with unequal variance, an unpaired t-test with Welch’s correction was used. One-way ANOVA followed by Tukey post hoc test was used for multiple group comparison. Significant difference is indicated by a p value less than 0.05 (*p < 0.05, **p < 0.01, ***p < 0.001, ****p < 0.0001). No statistical methods were used to pre-determine sample sizes but our sample sizes are similar to those reported in previous publications. Experiments were not randomized. Investigators were blinded as to the animal genotype during tissue section staining, image acquisition, and image analysis.

## Acknowledgements

We thank Dr. Junlei Chang for providing BAT-Gal mice. We thank the Core Facility and the Animal Facility of Medical Research Institute of Wuhan University for technical support. We thank all Zhou lab members for critical reading of the manuscript. Yan Zhou was supported by grants from National Key R&D Program of China (2022YFA0806603 and 2018YFA0800700), National Natural Science Foundation of China (31970770 and 32270876) and the Fundamental Research Funds for the Central Universities. Ying Liu was supported by grants from National Key R&D Program of China (2018YFA0800700 and 2022YFA0806603), National Natural Science Foundation of China (31970676) and Hubei Natural Science Foundation (2022CFB128).

## Supplementary figures

**Figure S1.**
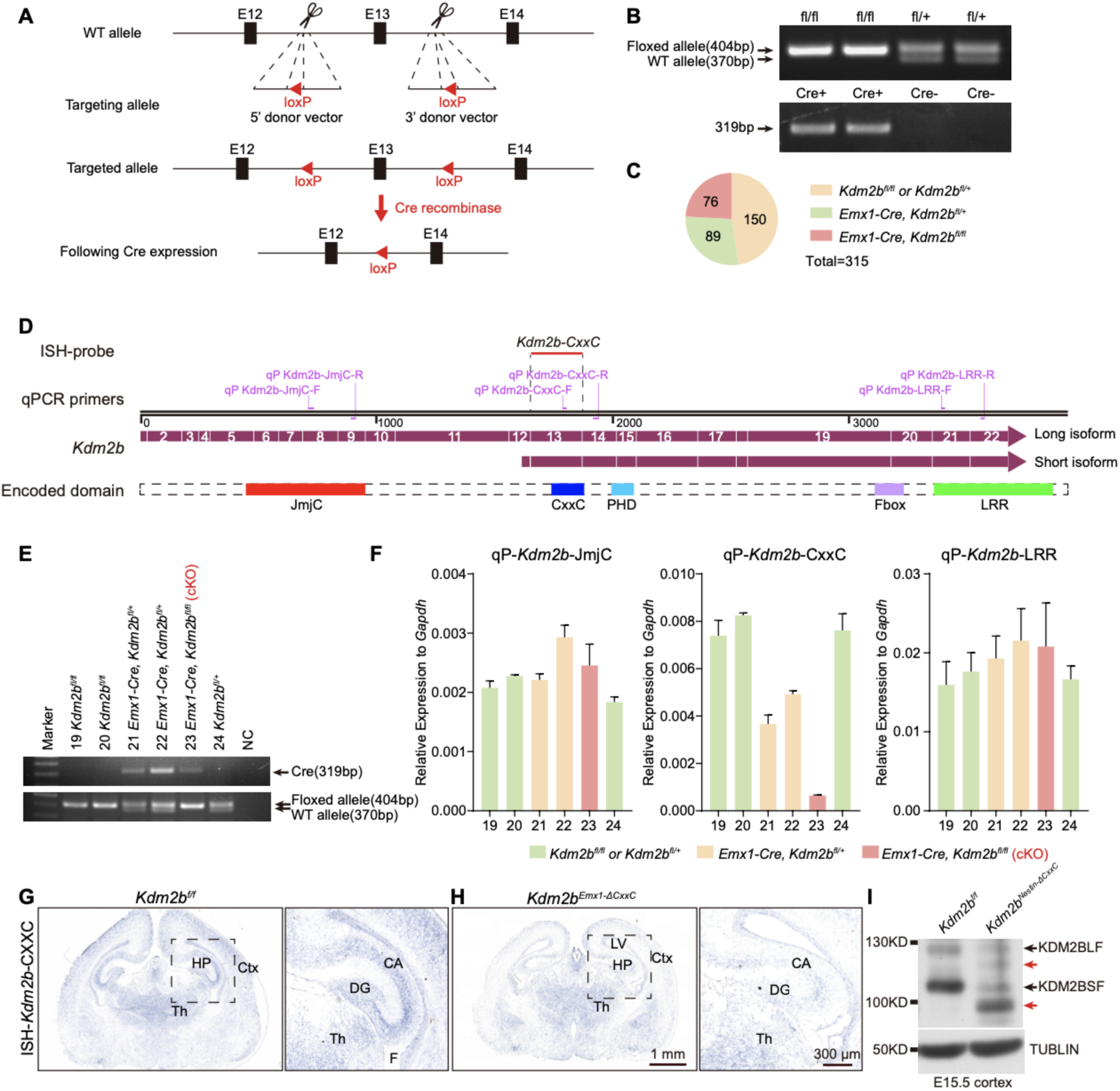
Selective ablation of the KDM2B-CxxC in the developing *Kdm2b^Emx1-ΔCxxC^* hippocampi. (A) Schematic representation of the *Kdm2b* genomic structure (top), targeting allele (middle) and targeted allele (bottom). Exon 13 is flanked by two loxP sites and will be excised after mating with Cre-recombinase-expressing mice. (B) PCR products for respective genotypes. (C) Offspring distribution of indicated genotypes at P0 from *Kdm2b^fl/fl^* females crossing with *Emx1-Cre*; *Kdm2b^fl/+^* males. (D) Schematic diagram of *Kdm2b* transcripts and corresponding protein domains. Positions of qRCR primers and the ISH probe were indicated. (E, F) Genotyping of a litter of P0 pups (**E**) and detection of *Kdm2b* expression by RT-qPCR (**F**). (G, H) Representative ISH images showing *Kdm2b* expression in coronal sections of P0 control (G) and *Kdm2b^Emx1-ΔCxxC^* (H) brains. Hippocampi (HP) were enlarged on the right. (I) Immunoblots of KDM2B and TUBLIN using extracts of E15.5 control and *Kdm2b^Nestin-ΔCxxC^* neocortices. Scale bars, 1mm (G, H left), 300 μm (G, H right). HP, Hippocampus; Ctx, cortex; Th, Thalamus; CA, Cornu Ammonis; DG, dentate gyrus; LV, lateral ventricle; F, fimbria.

**Figure supplement 1-source data**

Figure S1B source data 1: Genotyping of Cre.

Figure S1B source data 2: Genotyping of *floxed Kdm2b*.

Figure S1E source data: Genotyping of a litter of P0 pups for RT-qPCR.

Figure S1I source data: Immunoblotting of KDM2B and TUBLIN (E15.5 control and *Kdm2b^Nestin-ΔCxxC^* neocortices).

**Figure S2.**
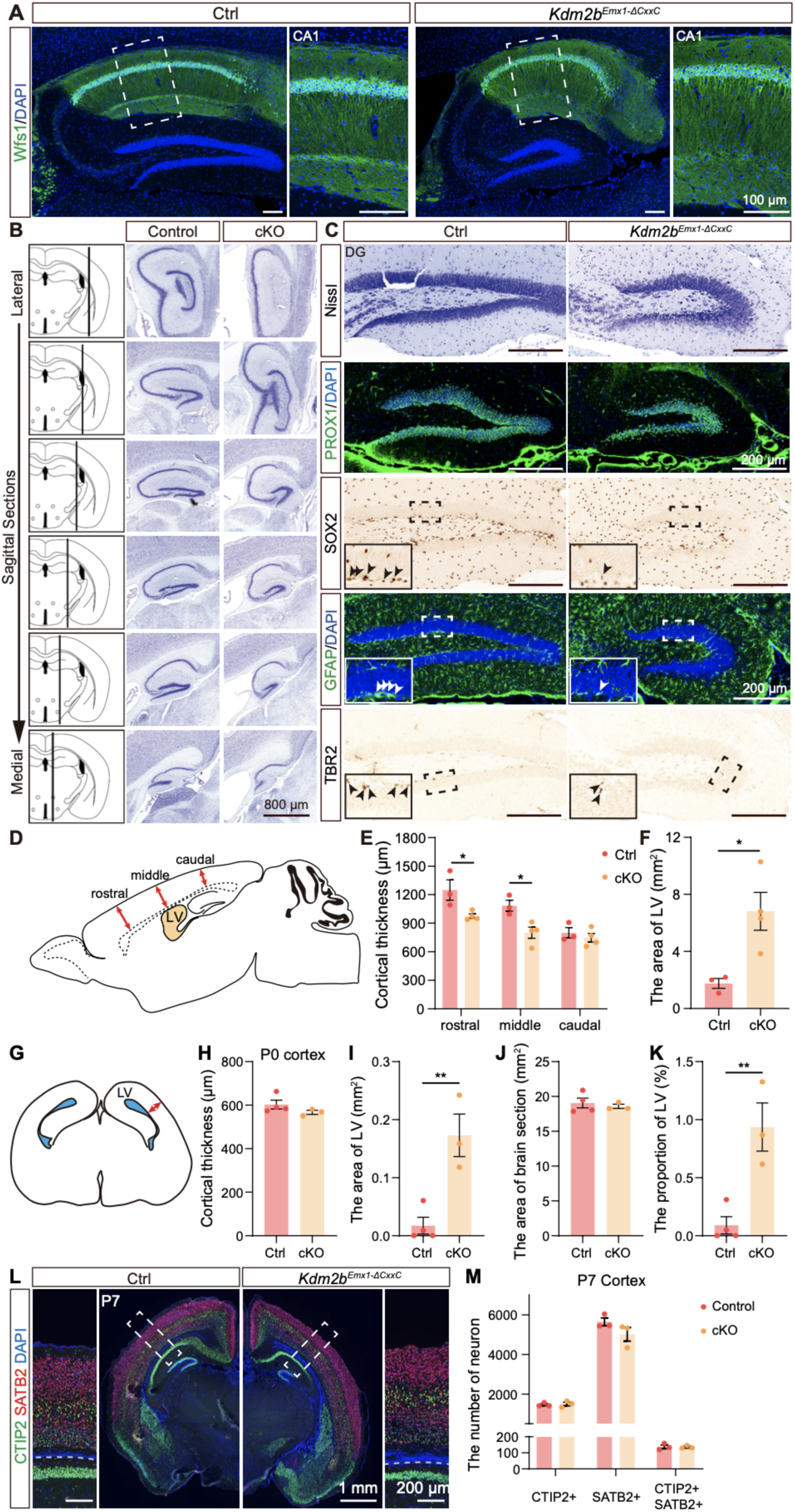
Deletion of the KDM2B-CxxC causes agenesis of hippocampus. (A) Immunofluorescent (IF) staining of Wfs1 on sagittal sections of adult control (left) and *Kdm2b^Emx1-ΔCxxC^* (right) hippocampi. Nuclei were labeled with DAPI (blue). Boxed regions of CA1 were enlarged on the right. (B) Nissl staining of sagittal sections of adult control and *Kdm2b^Emx1-ΔCxxC^* hippocampi. Sections from lateral to medial were displayed sequentially from top to bottom, with relative positions shown as black lines on the left. (C) From top to bottom: Nissl staining, IF staining for PROX1, IHC for SOX2, IF for GFAP and IHC for TBR2, on sagittal sections of adult control (left) and *Kdm2b^Emx1-ΔCxxC^* (right) DG. Boxed regions of SGZ were enlarged on the left-bottom corners. (D) The schematic diagram of a sagittal section of adult brain. Red bidirectional arrows show the rostral, middle, and caudal regions of the neocortex. (E) Quantifications of neocortical thickness of adult control and *Kdm2b^Emx1-ΔCxxC^* brains in rostral, middle, and caudal regions. (F) Quantifications of the area of lateral ventricle in adult control and *Kdm2b^Emx1-ΔCxxC^* brain. n = 3 for control brains and n = 4 for *Kdm2b^Emx1-ΔCxxC^* brains. (G) The diagram of a coronal section of P0 brain. The red bidirectional arrow indicates neocortical thickness. The blue areas indicate the lateral ventricles (LV). (H-K) Quantifications of neocortical thickness (H), area of lateral ventricles (LV) (I), area of brain sections (J) and the area proportion of LV (K) in P0 control and *Kdm2b^Emx1-ΔCxxC^* brain. n = 4 for control brains and n = 3 for *Kdm2b^Emx1-ΔCxxC^* brains. (L) IF staining for CTIP2 (green) and SATB2 (red) on coronal sections of P7 control (left) and *Kdm2b^Emx1-ΔCxxC^* (right) mice. Nuclei were labeled with DAPI (blue). Boxed regions were enlarged on the right. (M) Quantification of CTIP2+, SATB2+ and CTIP2+SATB2+ cells in P7 cortex (L). n = 3 for control brains and n = 3 for *Kdm2b^Emx1-ΔCxxC^* brains. Data are represented as means ± SEM. Statistical significance was determined using two-way ANOVA followed by Sidak’s multiple comparisons test (E), or using an unpaired two-tailed Student’s t-test (F). *P<0.05, **P < 0.01. Scale bars, 100 μm (A), 800 μm (B), 200 μm (C), 1 mm (L, whole brain), 200 μm (L, cortex). LV, lateral ventricles.

**Figure S3.**
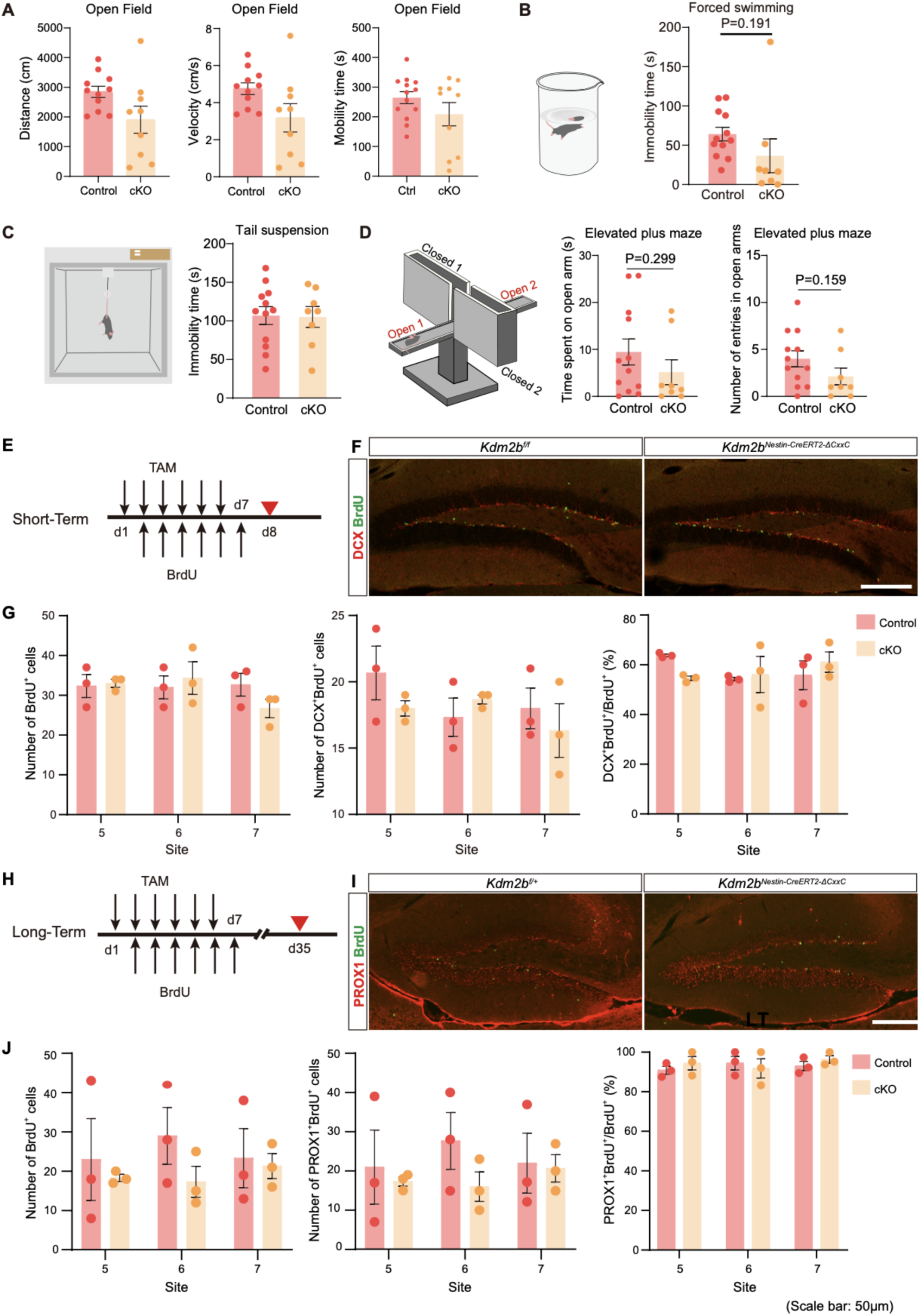
Adult neurogenesis in the SGZ was unaffected on conditional loss of KDM2B-CxxC. (A) Quantification of distance, velocity and mobility time in the open-field test. (B, C) Immobility time in forced swimming (B) and tail suspension experiments (C). (D) Time spent in open arms and the number of entries in open arms in elevated plus maze test. (E, H) Eight-week-old mice (*Kdm2b^f/f^* or *Kdm2b^f/+^*, and *Kdm2b^Nestin-CreERT2-ΔCxxC^*) were intraperitoneally injected with TAM and BrdU for 6 consecutive days (BrdU injection delayed by 1 day). Mice were sacrificed either 1 day later (Short-Term) or 4 weeks later (Long-Term). (F) Representative IF images showing DG sections stained with DCX (red) and BrdU (green). (G)Quantification of numbers of BrdU+ cells, DCX+BrdU+ cells, and the proportion of DCX+BrdU+/BrdU+ in the Short-Term experiment. (I) Representative IF images showing DG sections stained with PROX1 (red) and BrdU (green). (J) Quantification of numbers of BrdU+ cells, PROX1+BrdU+ cells, and the proportion of PROX1+BrdU+/BrdU+ in the Long-Term experiment. In (A), n = 11 mice for control and n = 9 mice for *Kdm2b^Emx1-ΔCxxC^*. In (B-D), n = 12 mice for control and n = 8 mice for *Kdm2b^Emx1-ΔCxxC^*. Data are represented as means ± SEM. Statistical significance was determined using an unpaired two-tailed Student’s t-test (A-D), or using two-way ANOVA followed by Sidak’s multiple comparisons test (G, J). In (E-J), n = 3 mice for both control and for *Kdm2b^Nestin-CreERT2-ΔCxxC^*. Scale bars, 50 μm (F, I).

**Figure S4.**
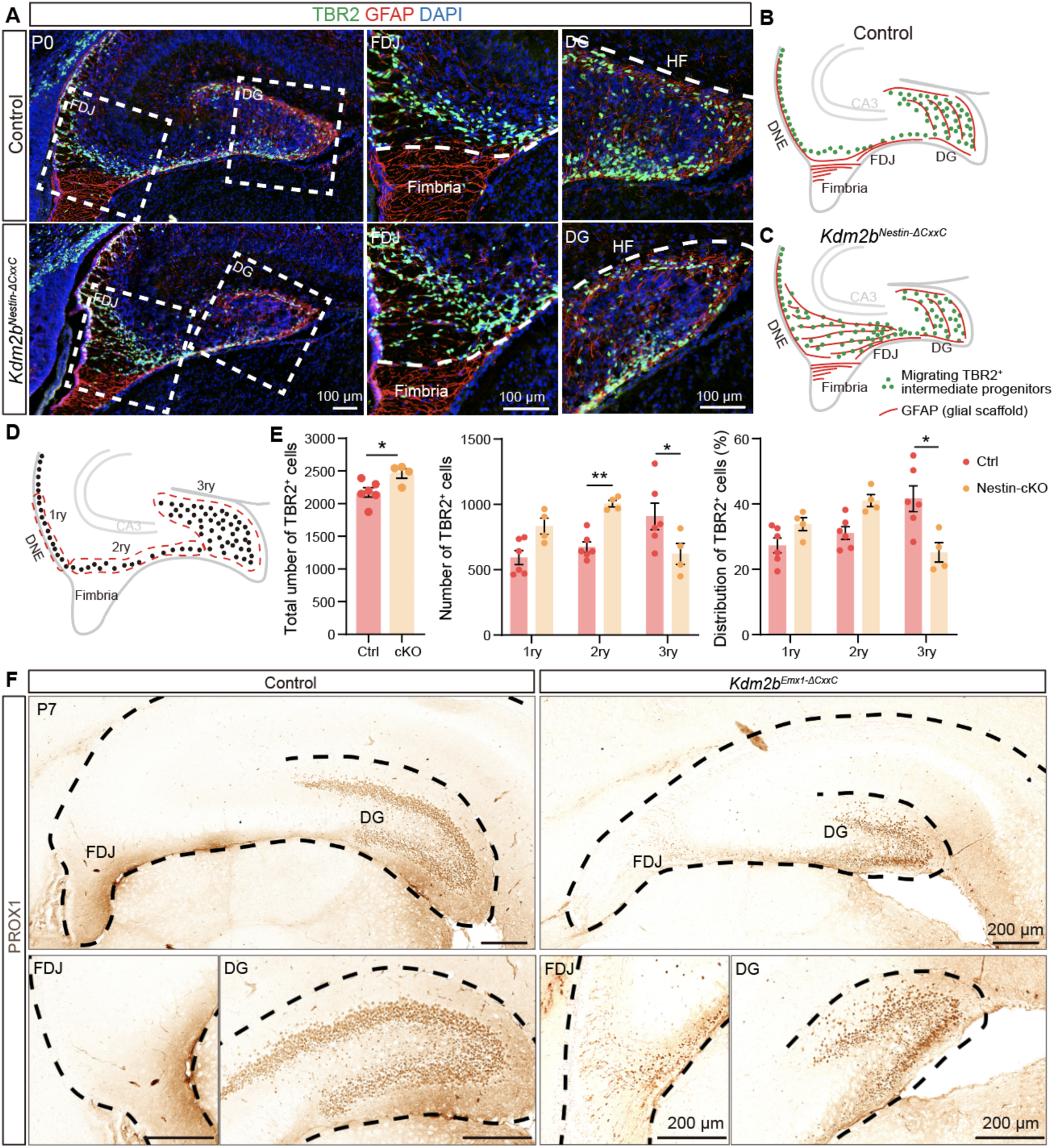
Ablation of the KDM2B-CxxC blocked the migration of intermediate progenitors and neurogenesis of granular cells. (A) Double immunofluorescence of TBR2 (green) and GFAP (red) on P0 wild-type and *Kdm2b^Nestin-ΔCxxC^* hippocampus. Nuclei were labeled with DAPI (blue). Boxed regions of FDJ and DG were enlarged on the right. (B, C) The schematic of P0 control and *Kdm2b^Nestin-ΔCxxC^* hippocampi. Green dots represent migrating TBR2+ intermediate progenitors, and red lines represent GFAP+ glial scaffold. (D, E) Distribution of TBR2+ cells along the three matrices, where dashed lines indicate areas considered as 1ry, 2ry, and 3ry matrix (D). (F) Immunohistochemical staining for PROX1 on coronal sections of P7 control (left) and *Kdm2b^Emx1-ΔCxxC^* (right) hippocampi. The DG and FDJ regions were individually enlarged underneath. n = 6 for control brains and n = 4 for *Kdm2b^Nestin-ΔCxxC^* brains. Data are represented as means ± SEM. Statistical significance was determined using an unpaired two-tailed Student’s t-test (E left), or using two-way ANOVA followed by Sidak’s multiple comparisons test (E middle and right). *P < 0.05, **P < 0.01. Scale bars, 100 μm (A), 200 μm (F). DG, dentate gyrus; DMS, dentate migratory stream; FDJ, fimbriodentate junction; HF, hippocampal fissure; 1ry, primary matrix; 2ry, secondary matrix; 3ry, tertiary matrix.

**Figure S5.**
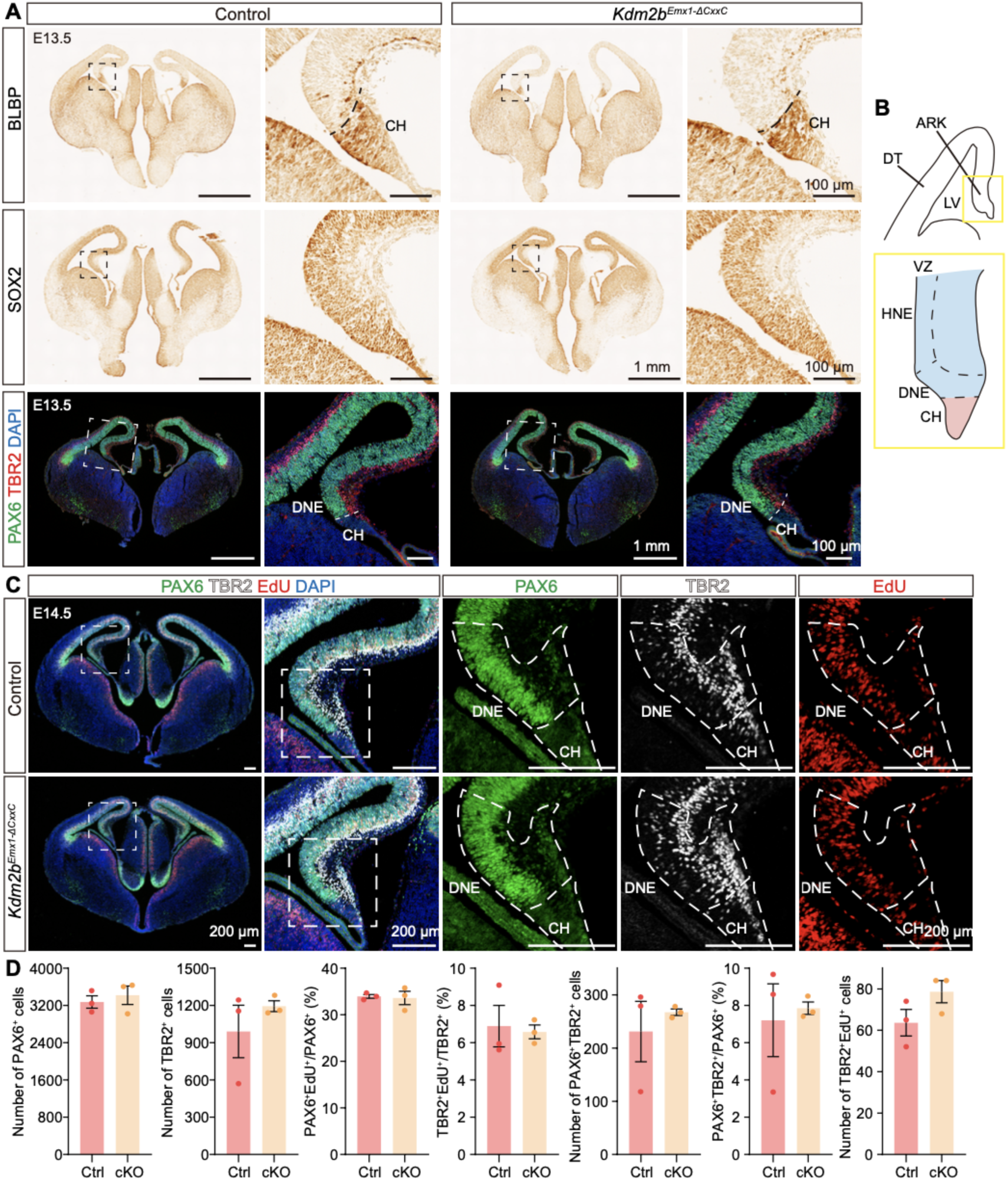
Unaltered neural progenitor domains in *Kdm2b^Emx1-ΔCxxC^* hippocampal primordia. (A) Immunohistochemical staining for BLBP (top) and SOX2 (middle). Bottom, immunofluorescent staining for PAX6 (green) and TBR2 (green) on coronal sections of E13.5 control (left) and *Kdm2b^Emx1-ΔCxxC^* (right) brain. Nuclei were labeled with DAPI (blue). Boxed regions were enlarged on the right. (B) The schematic of E13.5 wild-type hippocampus. The yellow box is enlarged to show the hippocampal primordia, with dashed lines demarcating HNE, DNE and CH. (C) Triple-labeling of PAX6 (green), TBR2 (white) and EdU (red) on E14.5 control and *Kdm2b^Emx1-ΔCxxC^* brain sections. Pregnant mice were injected with EdU 2 h before sacrifice. Nuclei were labeled with DAPI (blue). Boxed regions are enlarged on the right, and single channel fluorescence staining of PAX6, TBR2 and EdU were shown respectively. Dashed lines indicate DNE and CH. (D) Quantification of data in (C). Quantification of number of PAX6+, TBR2+, PAX6+TBR2+ and TBR2+EdU+ cells, and the proportion of PAX6+EdU+/PAX6+, TBR2+EdU+/TBR2+ and PAX6+TBR2+/PAX6+ in E14.5 control and *Kdm2b^Emx1-ΔCxxC^* DNE. n = 3 for control brains and n = 3 for *Kdm2b^Emx1-ΔCxxC^* brains. Data are represented as means ± SEM. Statistical significance was determined using an unpaired two-tailed Student’s t-test (D). Scale bars, 1 mm (A, whole brain), 100 μm (A, HP), 200 μm (C). CH, cortical hem; LV, lateral ventricle; DT, dorsal telencephalon; ARK, archicortex; HNE, hippocampal neuroepithelium; DNE, dentate neuroepithelium; VZ, ventricular zone.

**Figure S6.**
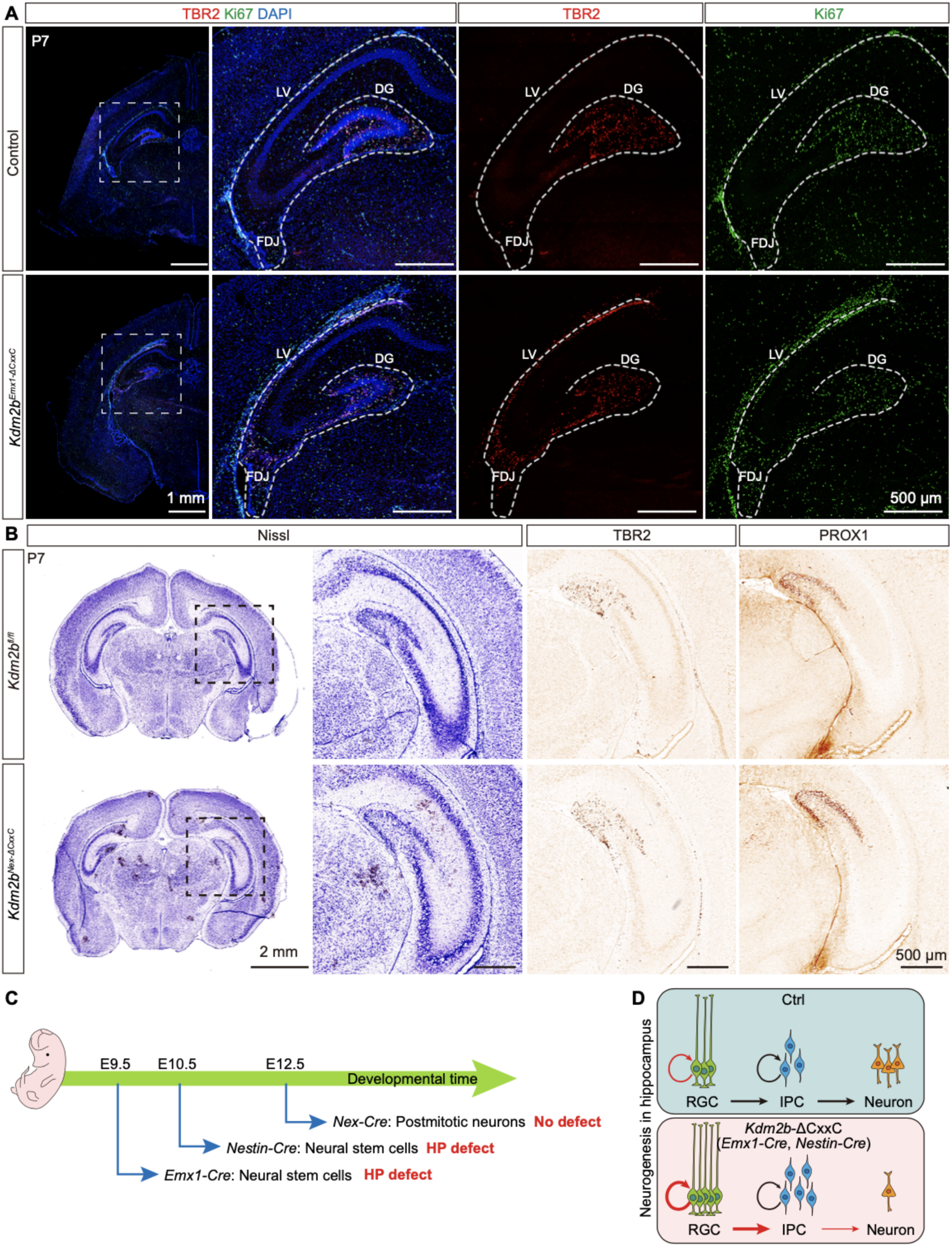
Neuronal differentiation is not responsible for hippocampal agenesis caused by loss of KDM2B-CxxC. (A) Double-labeling of TBR2 (red) and Ki67 (green) on P7 coronal section of control and *Kdm2b^Emx1-ΔCxxC^* brains. Nuclei were labeled with DAPI (blue). Boxed regions are enlarged on the right, and single channel fluorescence staining of TBR2 and Ki67 were shown respectively. Dashed lines outline the hippocampi. (B) Nissl staining, and immunohistochemical staining of TBR2 and PROX1 on coronal sections of P7 control and *Kdm2b^Nex-ΔCxxC^* brains. (C) The schematic diagram summarizing expression profiles of three Cre lines, as well as phenotypes of respective cKO mice. (D) The diagram showing aberrant neurogenesis during hippocampal development of *Kdm2b^Emx1-ΔCxxC^* and *Kdm2b^Nestin-ΔCxxC^* mice: a bigger RGC pool with more IPCs produced, but differentiation of IPCs toward neurons was compromised. Scale bars, 1 mm (A, whole brain), 100 μm (A, HP), 2 mm (B, whole brain), 500 μm (B, HP). LV, lateral ventricle; FDJ, fimbriodentate junction; DG, Dentate gyrus; RGC, Radial glia cells; IPC, Intermediate progenitor cell.

**Figure S7.**
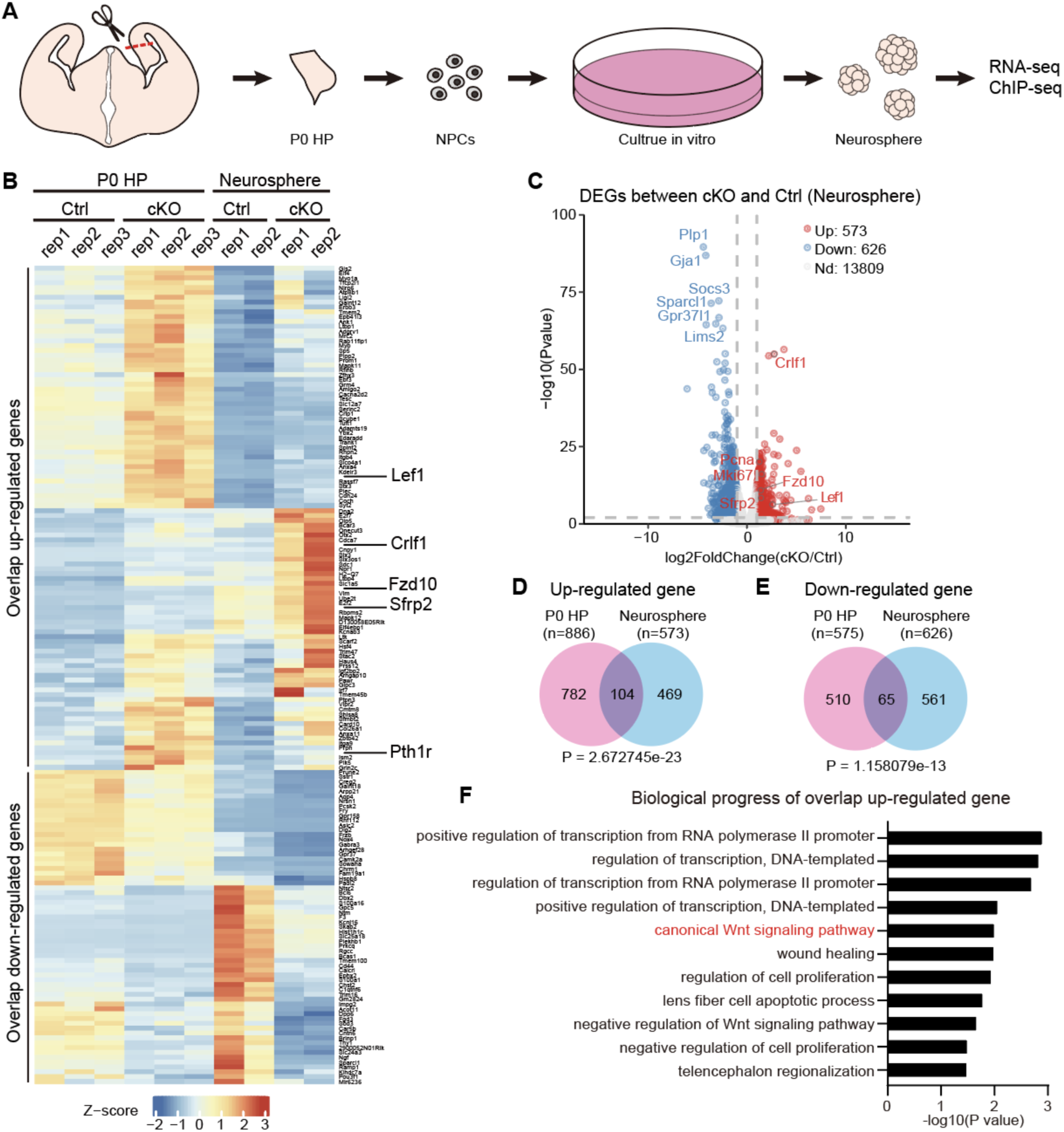
RNA-seq of neurospheres derived from P0 cKO hippocampi showed activation of the Wnt signaling pathway. (A) Schematic diagram: hippocampal tissue was collected from P0 brain and digested into single-cell suspensions. After cultured *in vitro* for three generations, neurospheres were subjected to RNA-seq and ChIP-seq analyses. (B) The heat map of overlapping up-and down-regulated genes of P0 HP and neurospheres. (C) The volcano plot of genes up-regulated (red) and down-regulated (blue) in P0 *Kdm2b^Emx1-ΔCxxC^* (cKO) neurospheres compared to controls. (D) Overlapping up-regulated genes (104) of P0 HP (782) and neurospheres (469). (E) Overlapping down-regulated genes (65) of P0 HP (510) and neurospheres (561). (F) GO analysis of the biological progress of overlapping up-regulated genes in *Kdm2b^Emx1-ΔCxxC^* (cKO) neurospheres revealed terms related to canonical Wnt signaling pathways (red).

**Figure S8.**
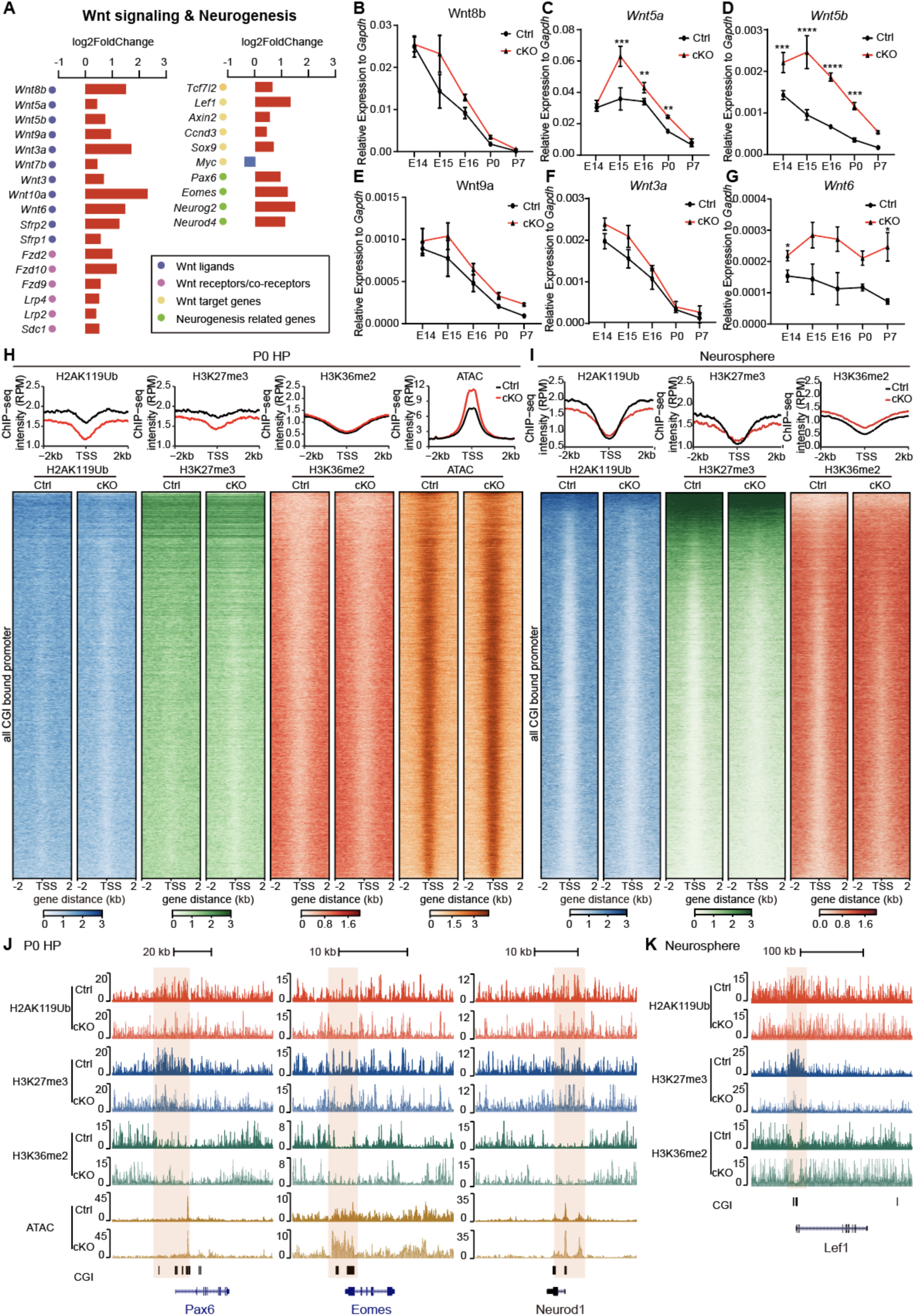
KDM2B epigenetically silences components of Wnt signaling genes in developing hippocampi. (A) Histograms showing the log-fold change of significantly up-or down-regulated genes in *Kdm2b^Emx1-ΔCxxC^* (cKO) hippocampi. (B-G) RT-qPCR showing relative expressions of *Wnt8b*, *Wnt5a*, *Wnt5b*, *Wnt9a*, *Wnt3a* and *Wnt6* in control (black lines) and *Kdm2b^Emx1-ΔCxxC^* (red lines) hippocampi of indicated developmental stages (E14.5, E15.5, E16.5, P0 and P7). (H) Heatmaps showing H2AK119ub, H3K27me3, H3K36me2, and ATAC-seq signals in control and *Kdm2b^Emx1-ΔCxxC^* (cKO) hippocampi at all CGI-associated promoters. Colors represent ChIP-seq RPM (reads per million), and rows were ranked by ChIP-seq signals in control H2AK119ub. Line charts on the top of each set of heatmap showing average signals, with control in black and *Kdm2b^Emx1-ΔCxxC^* (cKO) in red. (I) Line charts and heatmaps showing H2AK119ub, H3K27me3 and H3K36me2 signals in control and *Kdm2b^Emx1-ΔCxxC^* (cKO) neurospheres at all CGI-associated promoters. (J) The UCSC genome browser view of HA2K119ub, H3K27me3 and H3K36me2 enrichment and ATAC-seq signal in P0 control and *Kdm2b^Emx1-ΔCxxC^* (cKO) hippocampi at *Pax6*, *Eomes* and *Neurod1*. CGIs were shown as black columns on the bottom. Colored regions marked enrichment differences between control and cKO. (K) The UCSC genome browser view of HA2K119ub, H3K27me3 and H3K36me2 enrichment in control and *Kdm2b^Emx1-ΔCxxC^* (cKO) neurospheres at *Lef1*. CGIs were shown as black columns on the bottom. Colored regions marked enrichment differences between control and cKO. Data are represented as means ± SEM. Statistical significance was determined using two-way ANOVA followed by Sidak’s multiple comparisons test (B-G). *P < 0.05, **P < 0.01, ***P < 0.001, and ****P < 0.0001.

**Figure S9.**
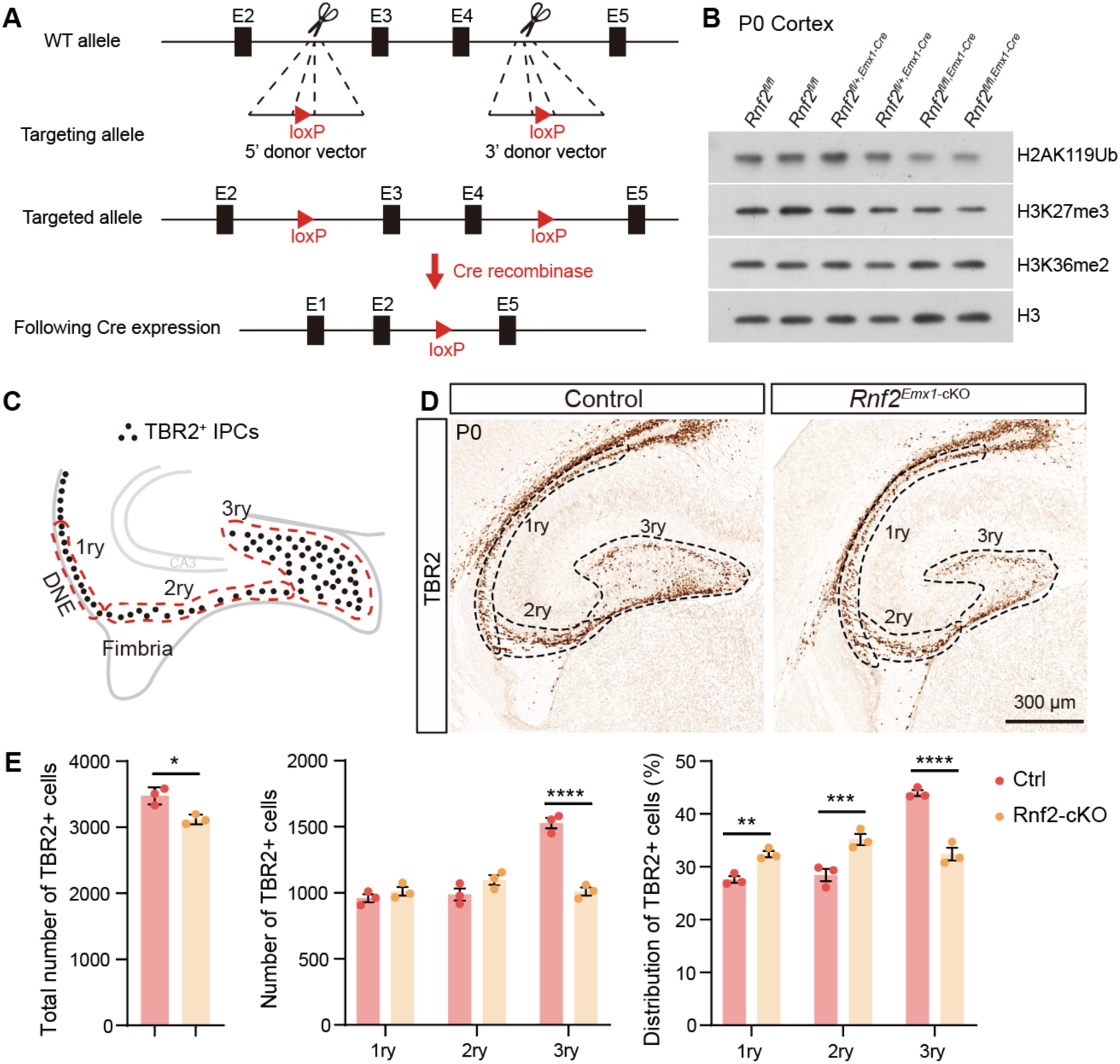
Loss of Ring1B did not cause accumulation of neural progenitors. (A) Schematic representation of the *Rnf2* genomic structure (top), targeting allele (middle) and targeted allele (bottom). Exon 3-4 is flanked by loxP sites and will be excised after mating with Cre-recombinase-expressing mice. (B) Immunoblots of H2AK119ub, H3K27me3, H3K36me2 and H3 using extracts of P0 Rnf2*^fl/fl^*, Rnf2*^fl/+, Emx1-Cre^* and Rnf2*^fl/fl, Emx1-Cre^* neocortices. (C-E) Immunohistochemical staining of TBR2 (D) on coronal sections of P0 control (left) and *Rnf2^Emx1-cKO^* (right) hippocampi. Distribution of TBR2+ cells along the three matrices, where dashed lines indicate areas considered as 1ry, 2ry, and 3ry matrix (C). n = 3 for control brains, n = 3 for *Rnf2 ^Emx1-cKO^* brains. Data are represented as means ± SEM. Statistical significance was determined using an unpaired two-tailed Student’s t-test (E, left) or using two-way ANOVA followed by Sidak’s multiple comparisons test (E, middle and right). *P < 0.05, **P < 0.01, ***P < 0.001, ****P < 0.0001, and “NS” indicates no significance. Scale bars, 300 μm (F). DNE, dentate neuroepithelium; 1ry, primary matrix; 2ry, secondary matrix; 3ry, tertiary matrix.

**Figure supplement 9-source data**

Figure S9B source data 1: Immunoblotting of H2AK119Ub (P0 *Rnf2^fl/fl^*, *Rnf2^fl/+, Emx1-Cre^* and Rnf2*^fl/fl, Emx1-Cre^* neocortices).

Figure S9B source data 2: Immunoblotting of H3K27me3 (P0 *Rnf2^fl/fl^*, *Rnf2^fl/+, Emx1-Cre^* and Rnf2*^fl/fl, Emx1-Cre^* neocortices).

Figure S9B source data 3: Immunoblotting of H3K36me2 (P0 *Rnf2^fl/fl^*, *Rnf2^fl/+, Emx1-Cre^* and Rnf2*^fl/fl, Emx1-Cre^* neocortices).

Figure S9B source data 4: Immunoblotting of H3 (P0 *Rnf2^fl/fl^*, *Rnf2^fl/+, Emx1-Cre^* and *Rnf2^fl/fl, Emx1-Cre^* neocortices).

## Supplemental Tables

**Supplemental Table S1.**
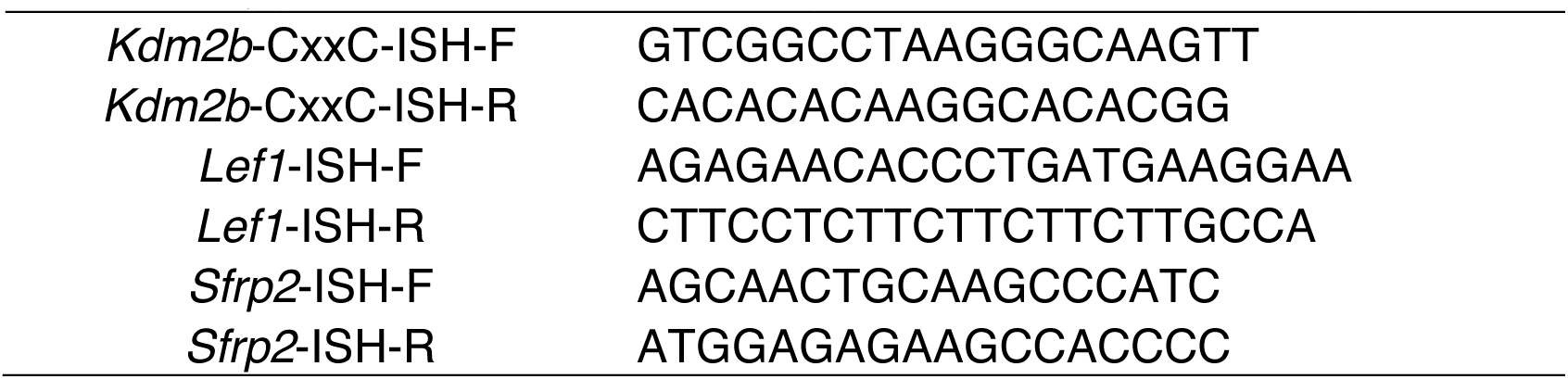
ISH primers used in this study

**Supplemental Table S2.**
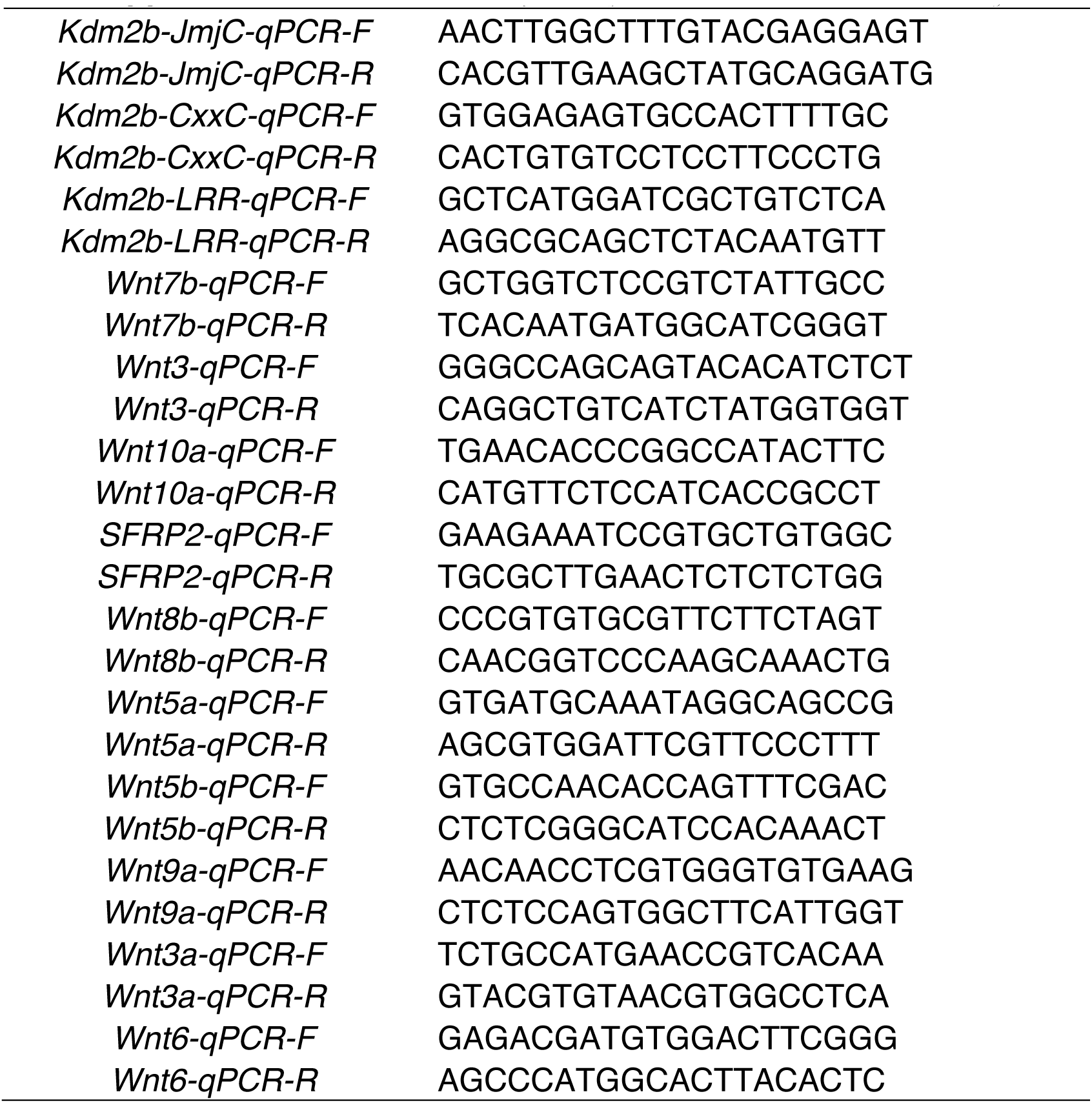
RT-qPCR primers used in this study

**Supplemental Table S3.**
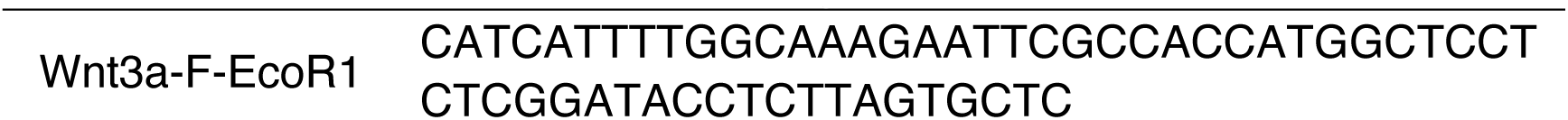
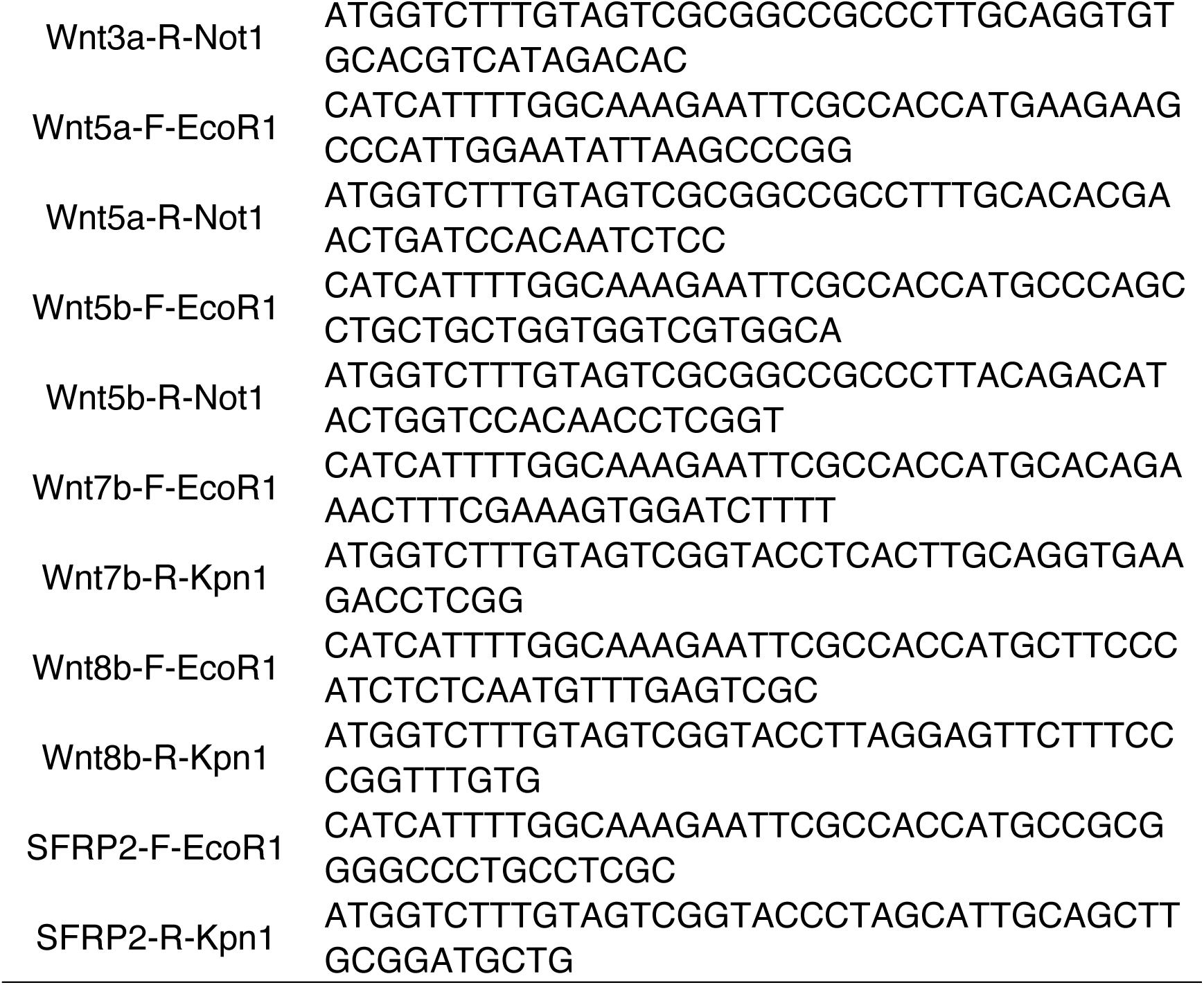
Plasmid construction primers used in this study

## References

1. Anacker, C., & Hen, R. (2017). Adult hippocampal neurogenesis and cognitive flexibility - linking memory and mood. Nat Rev Neurosci, 18(6), 335–346. doi:10.1038/nrn.2017.45

2. Bengoa-Vergniory, Nora, & Kypta, Robert M. (2015). Canonical and noncanonical Wnt signaling in neural stem/progenitor cells. Cellular and Molecular Life Sciences, 72(21), 4157–4172. doi:10.1007/s00018-015-2028-6

3. Bingman, Verner P., Salas, Cosme, & Rodriguez, Fernando. (2009). Evolution of the Hippocampus. In Marc D. Binder, Nobutaka Hirokawa, & Uwe Windhorst (Eds.), Encyclopedia of Neuroscience (pp. 1356–1360). Berlin, Heidelberg: Springer Berlin Heidelberg.

4. Blackledge, N. P., Farcas, A. M., Kondo, T., King, H. W., McGouran, J. F., Hanssen, L. L., … Klose, R. J. (2014). Variant PRC1 complex-dependent H2A ubiquitylation drives PRC2 recruitment and polycomb domain formation. Cell, 157(6), 1445–1459. doi:10.1016/j.cell.2014.05.004

5. Blackledge, N. P., & Klose, R. J. (2021). The molecular principles of gene regulation by Polycomb repressive complexes. Nat Rev Mol Cell Biol, 22(12), 815–833. doi:10.1038/s41580-021-00398-y

6. Bonnet, J., Boichenko, I., Kalb, R., Le Jeune, M., Maltseva, S., Pieropan, M., … Muller, J. (2022). PR-DUB preserves Polycomb repression by preventing excessive accumulation of H2Aub1, an antagonist of chromatin compaction. Genes Dev, 36(19-20), 1046–1061. doi:10.1101/gad.350014.122

7. Caramello, A., Galichet, C., Rizzoti, K., & Lovell-Badge, R. (2021). Dentate gyrus development requires a cortical hem-derived astrocytic scaffold. elife, 10. doi:10.7554/eLife.63904

8. Chen, Z., Djekidel, M. N., & Zhang, Y. (2021). Distinct dynamics and functions of H2AK119ub1 and H3K27me3 in mouse preimplantation embryos. Nat Genet, 53(4), 551–563. doi:10.1038/s41588-021-00821-2

9. Chenn, A., & Walsh, C. A. (2002). Regulation of cerebral cortical size by control of cell cycle exit in neural precursors. Science, 297(5580), 365–369. doi:10.1126/science.1074192

10. Chiacchiera, Fulvio, Rossi, Alessandra, Jammula, SriGanesh, Piunti, Andrea, Scelfo, Andrea, Ordóñez-Morán, Paloma, … Pasini, Diego. (2016). Polycomb Complex PRC1 Preserves Intestinal Stem Cell Identity by Sustaining Wnt/β-Catenin Transcriptional Activity. Cell Stem Cell, 18(1), 91–103. doi:https://doi.org/10.1016/j.stem.2015.09.019

11. Eto, H., Kishi, Y., Yakushiji-Kaminatsui, N., Sugishita, H., Utsunomiya, S., Koseki, H., & Gotoh, Y. (2020). The Polycomb group protein Ring1 regulates dorsoventral patterning of the mouse telencephalon. Nat Commun, 11(1), 5709. doi:10.1038/s41467-020-19556-5

12. Farcas, A. M., Blackledge, N. P., Sudbery, I., Long, H. K., McGouran, J. F., Rose, N. R., … Klose, R. J. (2012). KDM2B links the Polycomb Repressive Complex 1 (PRC1) to recognition of CpG islands. Elife, 1, e00205. doi:10.7554/eLife.00205

13. Fukuda, T., Tokunaga, A., Sakamoto, R., & Yoshida, N. (2011). Fbxl10/Kdm2b deficiency accelerates neural progenitor cell death and leads to exencephaly. Mol Cell Neurosci, 46(3), 614–624. doi:10.1016/j.mcn.2011.01.001

14. Fursova, N. A., Blackledge, N. P., Nakayama, M., Ito, S., Koseki, Y., Farcas, A. M., … Klose, R. J. (2019). Synergy between Variant PRC1 Complexes Defines Polycomb-Mediated Gene Repression. Mol Cell, 74(5), 1020–1036 e1028. doi:10.1016/j.molcel.2019.03.024

15. Galceran, J., Miyashita-Lin, E.M., Devaney, E., Rubenstein, J.L., & Grosschedl, R. (2000). Hippocampus development and generation of dentate gyrus granule cells is regulated by LEF1. Development, 127(3), 469–482. doi:10.1242/dev.127.3.469

16. Gao, Yuen, Duque-Wilckens, Natalia, Aljazi, Mohammad B, Moeser, Adam J, Mias, George I, Robison, Alfred J, … He, Jin. (2022). Impaired KDM2B-mediated PRC1 recruitment to chromatin causes defective neural stem cell self-renewal and ASD/ID-like behaviors. iScience, 103742.

17. Gao, Z., Zhang, J., Bonasio, R., Strino, F., Sawai, A., Parisi, F., … Reinberg, D. (2012). PCGF homologs, CBX proteins, and RYBP define functionally distinct PRC1 family complexes. Mol Cell, 45(3), 344–356. doi:10.1016/j.molcel.2012.01.002

18. Gearhart, M. D., Corcoran, C. M., Wamstad, J. A., & Bardwell, V. J. (2006). Polycomb group and SCF ubiquitin ligases are found in a novel BCOR complex that is recruited to BCL6 targets. Mol Cell Biol, 26(18), 6880–6889. doi:10.1128/MCB.00630-06

19. Guan, H., Zhang, J., Luan, J., Xu, H., Huang, Z., Yu, Q., … Xu, L. (2021). Secreted Frizzled Related Proteins in Cardiovascular and Metabolic Diseases. Front Endocrinol (Lausanne), 12, 712217. doi:10.3389/fendo.2021.712217

20. He, J., Shen, L., Wan, M., Taranova, O., Wu, H., & Zhang, Y. (2013). Kdm2b maintains murine embryonic stem cell status by recruiting PRC1 complex to CpG islands of developmental genes. Nat Cell Biol, 15(4), 373–384. doi:10.1038/ncb2702

21. Hirabayashi, Y., Suzki, N., Tsuboi, M., Endo, T. A., Toyoda, T., Shinga, J., … Gotoh, Y. (2009). Polycomb limits the neurogenic competence of neural precursor cells to promote astrogenic fate transition. Neuron, 63(5), 600–613. doi:10.1016/j.neuron.2009.08.021

22. Huo, D., Yu, Z., Li, R., Gong, M., Sidoli, S., Lu, X., … Wu, X. (2022). CpG island reconfiguration for the establishment and synchronization of polycomb functions upon exit from naive pluripotency. Mol Cell. doi:10.1016/j.molcel.2022.01.027

23. Kalb, R., Latwiel, S., Baymaz, H. I., Jansen, P. W., Muller, C. W., Vermeulen, M., & Muller, J. (2014). Histone H2A monoubiquitination promotes histone H3 methylation in Polycomb repression. Nat Struct Mol Biol, 21(6), 569–571. doi:10.1038/nsmb.2833

24. Labonne, J. D., Lee, K. H., Iwase, S., Kong, I. K., Diamond, M. P., Layman, L. C., … Kim, H.G. (2016). An atypical 12q24.31 microdeletion implicates six genes including a histone demethylase KDM2B and a histone methyltransferase SETD1B in syndromic intellectual disability. Hum Genet, 135(7), 757–771. doi:10.1007/s00439-016-1668-4

25. Lan, Xianchun, Ding, Song, Zhang, Tianzhe, Yi, Ying, Li, Conghui, Jin, Wenwen, … Jiang, Wei. (2022). PCGF6 controls neuroectoderm specification of human pluripotent stem cells by activating SOX2 expression. Nature Communications, 13(1), 4601. doi:10.1038/s41467-022-32295-z

26. Laugesen, A., & Helin, K. (2014). Chromatin repressive complexes in stem cells, development, and cancer. Cell Stem Cell, 14(6), 735–751. doi:10.1016/j.stem.2014.05.006

27. Li, G., & Pleasure, S. J. (2014). The development of hippocampal cellular assemblies. Wiley Interdiscip Rev Dev Biol, 3(2), 165–177. doi:10.1002/wdev.127

28. Li, W., Shen, W., Zhang, B., Tian, K., Li, Y., Mu, L., … Zhou, Y. (2020). Long non-coding RNA LncKdm2b regulates cortical neuronal differentiation by cis-activating Kdm2b. Protein Cell, 11(3), 161–186. doi:10.1007/s13238-019-0650-z

29. Lie, Dieter-Chichung, Colamarino, Sophia A., Song, Hong-Jun, Désiré, Laurent, Mira, Helena, Consiglio, Antonella, … Gage, Fred H. (2005). Wnt signalling regulates adult hippocampal neurogenesis. Nature, 437(7063), 1370–1375. doi:10.1038/nature04108

30. Liu, P. P., Xu, Y. J., Dai, S. K., Du, H. Z., Wang, Y. Y., Li, X. G., … Liu, C. M. (2019). Polycomb Protein EED Regulates Neuronal Differentiation through Targeting SOX11 in Hippocampal Dentate Gyrus. Stem Cell Reports, 13(1), 115–131. doi:10.1016/j.stemcr.2019.05.010

31. Machon, O., van den Bout, C. J., Backman, M., Kemler, R., & Krauss, S. (2003). Role of β- catenin in the developing cortical and hippocampal neuroepithelium. Neuroscience, 122(1), 129–143. doi:https://doi.org/10.1016/S0306-4522(03)00519-0

32. Marcos, S., Nieto-Lopez, F., Sandonis, A., Cardozo, M. J., Di Marco, F., Esteve, P., & Bovolenta, P. (2015). Secreted frizzled related proteins modulate pathfinding and fasciculation of mouse retina ganglion cell axons by direct and indirect mechanisms. J Neurosci, 35(11), 4729–4740. doi:10.1523/JNEUROSCI.3304-13.2015

33. Margueron, Raphael, Justin, Neil, Ohno, Katsuhito, Sharpe, Miriam L., Son, Jinsook, Drury Iii, William J., … Gamblin, Steven J. (2009). Role of the polycomb protein EED in the propagation of repressive histone marks. Nature, 461(7265), 762–767. doi:10.1038/nature08398

34. Morimoto-Suzki, N., Hirabayashi, Y., Tyssowski, K., Shinga, J., Vidal, M., Koseki, H., & Gotoh, Y. (2014). The polycomb component Ring1B regulates the timed termination of subcerebral projection neuron production during mouse neocortical development. Development, 141(22), 4343–4353. doi:10.1242/dev.112276

35. Munji, R. N., Choe, Y., Li, G., Siegenthaler, J. A., & Pleasure, S. J. (2011). Wnt signaling regulates neuronal differentiation of cortical intermediate progenitors. J Neurosci, 31(5), 1676–1687. doi:10.1523/JNEUROSCI.5404-10.2011

36. Nelson, Branden R., Hodge, Rebecca D., Daza, Ray A. M., Tripathi, Prem Prakash, Arnold, Sebastian J., Millen, Kathleen J., & Hevner, Robert F. (2020). Intermediate progenitors support migration of neural stem cells into dentate gyrus outer neurogenic niches. elife, 9, e53777. doi:10.7554/eLife.53777

37. O’Carroll, D., Erhardt, S., Pagani, M., Barton, S. C., Surani, M. A., & Jenuwein, T. (2001). The polycomb-group gene Ezh2 is required for early mouse development. Mol Cell Biol, 21(13), 4330–4336. doi:10.1128/mcb.21.13.4330-4336.2001

38. Oittinen, Mikko, Popp, Alina, Kurppa, Kalle, Lindfors, Katri, Mäki, Markku, Kaikkonen, Minna U., & Viiri, Keijo. (2016). Polycomb Repressive Complex 2 Enacts Wnt Signaling in Intestinal Homeostasis and Contributes to the Instigation of Stemness in Diseases Entailing Epithelial Hyperplasia or Neoplasia. Stem Cells, 35(2), 445–457. doi:10.1002/stem.2479

39. Pereira, J. D., Sansom, S. N., Smith, J., Dobenecker, M. W., Tarakhovsky, A., & Livesey, F. J. (2010). Ezh2, the histone methyltransferase of PRC2, regulates the balance between self-renewal and differentiation in the cerebral cortex. Proc Natl Acad Sci U S A, 107(36), 15957–15962. doi:10.1073/pnas.1002530107

40. Piper, M., Barry, G., Harvey, T. J., McLeay, R., Smith, A. G., Harris, L., … Richards, L. J. (2014). NFIB-mediated repression of the epigenetic factor Ezh2 regulates cortical development. J Neurosci, 34(8), 2921–2930. doi:10.1523/JNEUROSCI.2319-13.2014

41. Piunti, A., & Shilatifard, A. (2016). Epigenetic balance of gene expression by Polycomb and COMPASS families. Science, 352(6290), aad9780. doi:10.1126/science.aad9780

42. Satoh, W., Gotoh, T., Tsunematsu, Y., Aizawa, S., & Shimono, A. (2006). Sfrp1 and Sfrp2 regulate anteroposterior axis elongation and somite segmentation during mouse embryogenesis. Development, 133(6), 989–999. doi:10.1242/dev.02274

43. Schuettengruber, B., & Cavalli, G. (2009). Recruitment of polycomb group complexes and their role in the dynamic regulation of cell fate choice. Development, 136(21), 3531–3542. doi:10.1242/dev.033902

44. Sugishita, H., Kondo, T., Ito, S., Nakayama, M., Yakushiji-Kaminatsui, N., Kawakami, E., … Koseki, H. (2021). Variant PCGF1-PRC1 links PRC2 recruitment with differentiation-associated transcriptional inactivation at target genes. Nat Commun, 12(1), 5341. doi:10.1038/s41467-021-24894-z

45. Sun, B., Chang, E., Gerhartl, A., & Szele, F. G. (2018). Polycomb Protein Eed is Required for Neurogenesis and Cortical Injury Activation in the Subventricular Zone. Cereb Cortex, 28(4), 1369–1382. doi:10.1093/cercor/bhx289

46. Supèr, H., Soriano, E., & Uylings, H. B. M. (1998). The functions of the preplate in development and evolution of the neocortex and hippocampus. Brain Research Reviews, 27(1), 40–64. doi:https://doi.org/10.1016/S0165-0173(98)00005-8

47. Tamburri, S., Lavarone, E., Fernandez-Perez, D., Conway, E., Zanotti, M., Manganaro, D., & Pasini, D. (2020). Histone H2AK119 Mono-Ubiquitination Is Essential for Polycomb-Mediated Transcriptional Repression. Mol Cell, 77(4), 840–856 e845. doi:10.1016/j.molcel.2019.11.021

48. Turberfield, A. H., Kondo, T., Nakayama, M., Koseki, Y., King, H. W., Koseki, H., & Klose, R. J. (2019). KDM2 proteins constrain transcription from CpG island gene promoters independently of their histone demethylase activity. Nucleic Acids Res, 47(17), 9005–9023. doi:10.1093/nar/gkz607

49. van Jaarsveld, R. H., Reilly, J., Cornips, M. C., Hadders, M. A., Agolini, E., Ahimaz, P., … Oegema, R. (2022). Delineation of a KDM2B-related neurodevelopmental disorder and its associated DNA methylation signature. Genet Med. doi:10.1016/j.gim.2022.09.006

50. von Marschall, Z., & Fisher, L. W. (2010). Secreted Frizzled-related protein-2 (sFRP2) augments canonical Wnt3a-induced signaling. Biochem Biophys Res Commun, 400(3), 299–304. doi:10.1016/j.bbrc.2010.08.043

51. von Schimmelmann, M., Feinberg, P. A., Sullivan, J. M., Ku, S. M., Badimon, A., Duff, M. K., … Schaefer, A. (2016). Polycomb repressive complex 2 (PRC2) silences genes responsible for neurodegeneration. Nat Neurosci, 19(10), 1321–1330. doi:10.1038/nn.4360

52. Voncken, J. W., Roelen, B. A., Roefs, M., de Vries, S., Verhoeven, E., Marino, S., … van Lohuizen, M. (2003). Rnf2 (Ring1b) deficiency causes gastrulation arrest and cell cycle inhibition. Proc Natl Acad Sci U S A, 100(5), 2468–2473. doi:10.1073/pnas.0434312100

53. Vorhees, C. V., & Williams, M. T. (2006). Morris water maze: procedures for assessing spatial and related forms of learning and memory. Nat Protoc, 1(2), 848–858. doi:10.1038/nprot.2006.116

54. Wang, Jiajia, Yang, Lijun, Dong, Chen, Wang, Jincheng, Xu, Lingli, Qiu, Yueping, … Lu, Q. Richard. (2020). EED-mediated histone methylation is critical for CNS myelination and remyelination by inhibiting WNT, BMP, and senescence pathways. Science Advances, 6(33), eaaz6477. doi:10.1126/sciadv.aaz6477

55. Wang, Junbao, Wang, Andi, Tian, Kuan, Hua, Xiaojiao, Zhang, Bo, Zheng, Yue, … Zhou, Yan. (2022). A Ctnnb1 enhancer regulates neocortical neurogenesis by controlling the abundance of intermediate progenitors. Cell Discovery, 8(1), 74. doi:10.1038/s41421-022-00421-2

56. Wang, Z., Gearhart, M. D., Lee, Y. W., Kumar, I., Ramazanov, B., Zhang, Y., … Ivanova, N. B. (2018). A Non-canonical BCOR-PRC1.1 Complex Represses Differentiation Programs in Human ESCs. Cell Stem Cell, 22(2), 235–251 e239. doi:10.1016/j.stem.2017.12.002

57. Wu, X., Johansen, J. V., & Helin, K. (2013). Fbxl10/Kdm2b Recruits Polycomb Repressive Complex 1 to CpG Islands and Regulates H2A Ubiquitylation. Mol Cell. doi:10.1016/j.molcel.2013.01.016

58. Yokotsuka-Ishida, S., Nakamura, M., Tomiyasu, Y., Nagai, M., Kato, Y., Tomiyasu, A., … Sano, A. (2021). Positional cloning and comprehensive mutation analysis identified a novel KDM2B mutation in a Japanese family with minor malformations, intellectual disability, and schizophrenia. J Hum Genet, 66(6), 597–606. doi:10.1038/s10038-020-00889-4

59. Zhang, J., Ji, F., Liu, Y., Lei, X., Li, H., Ji, G., … Jiao, J. (2014). Ezh2 regulates adult hippocampal neurogenesis and memory. J Neurosci, 34(15), 5184–5199. doi:10.1523/JNEUROSCI.4129-13.2014

60. Zhong, S., Ding, W., Sun, L., Lu, Y., Dong, H., Fan, X., … Wang, X. (2020). Decoding the development of the human hippocampus. Nature, 577(7791), 531–536. doi:10.1038/s41586-019-1917-5

